# Cyclic-di-adenosine monophosphate (c-di-AMP) is required for osmotic regulation in *Staphylococcus aureus* but dispensable for viability in anaerobic conditions

**DOI:** 10.1101/216887

**Authors:** Merve S. Zeden, Christopher F. Schuster, Lisa Bowman, Qiyun Zhong, Huw D. Williams, Angelika Gründling

## Abstract

Cyclic di-adenosine monophosphate (c-di-AMP) is a recently discovered signaling molecule important for the survival of Firmicutes, a large bacterial group that includes notable pathogens such as *Staphylococcus aureus*. However, the exact role of this molecule has not been identified. *dacA*, the *S. aureus* gene encoding the diadenylate cyclase enzyme required for c-di-AMP production, cannot be deleted when bacterial cells are grown in rich medium, indicating that c-di-AMP is required for growth in this condition. Here, we report that an *S. aureus dacA* mutant can be generated in chemically defined medium. Consistent with previous findings, this mutant had a severe growth defect when cultured in rich medium. Using this growth defect in rich medium, we selected for suppressor strains with improved growth to identify c-di-AMP-requiring pathways. Mutations bypassing the essentiality of *dacA* were identified in *alsT* and *opuD*, encoding a predicted amino acid and osmolyte transporter, the latter of which we show here to be the main glycine betaine-uptake system in *S. aureus*. Inactivation of these transporters likely prevents the excessive osmolyte and amino acid accumulation in the cell, providing further evidence for a key role of c-di-AMP in osmotic regulation. Suppressor mutations were also obtained in *hepS, hemB, ctaA* and *qoxB*, coding for proteins required for respiration. Furthermore, we show that *dacA* is dispensable for growth in anaerobic conditions. Together, these finding reveal an essential role for the c-di-AMP signaling network in aerobic, but not anaerobic, respiration in *S. aureus*.

---

*Staphylococcus aureus* colonizes the nares and skin of humans permanently or transiently and can cause infections ranging from skin and soft tissue infections to endocarditis and bacteremia (1,2). Infections of *S. aureus* are becoming more difficult to treat due to the emergence of antibiotic resistant strains such as methicillin resistant *S. aureus* (MRSA) strains. In addition, the increased prevalence of community-acquired methicillin resistant *S. aureus* (CA-MRSA) strains in parts of the world pose a significant threat to public health (1–3). During the course of an infection, oxygen availability is an environmental factor that can vary dramatically and when *S. aureus* migrates from the nasal cavity or the skin to internal tissues, the availability of free oxygen decreases significantly (4). In order to proliferate under oxygen limiting conditions, *S. aureus* shifts its metabolism and generates energy by for instance anaerobic nitrate respiration or through fermentation (4–6).

When *S. aureus* respires aerobically, various organic substrates, including NADH, succinate and lactate, are oxidized by their corresponding dehydrogenase enzymes and the freed electrons are passed to terminal oxidases via menaquinones (MQ) and ultimately used to reduce O_2_ to H_2_O (7-9). During this process, protons are extruded and the proton motive force that is generated is used for the production of ATP via the F_1_F_0_-ATPase (9). *S. aureus* has a branched aerobic respiratory chain and possesses two heme-dependent terminal oxidases referred to as cytochrome aa3 (cyt aa3) and cytochrome bd (cyt bd) (10). They are also referred to as the Qox system and the Cyd system and the former is the main terminal oxidase used by *S. aureus* during aerobic respiration (10). *S. aureus* can also respire under anaerobic conditions using nitrate as a terminal electron acceptor and generate the membrane potential in this manner. In the absence of a suitable electron acceptor, *S. aureus* will grow by fermentation under anaerobic conditions (6).

Secondary messenger signaling networks are frequently utilized by bacteria to rapidly adapt to changing environments (11,12). The regulatory actions of secondary messengers can range from transcriptional to post-transcriptional and even post-translational mechanisms (12-14). Cyclic diadenosine monophosphate (c-di-AMP) is a recently discovered secondary messenger molecule that is produced predominantly by Gram-positive bacteria (11,13,15-18). In *S. aureus*, c-di-AMP is synthesized from two molecules of ATP by the diadenylate cyclase enzyme DacA and degraded into pApA by the phosphodiesterase (PDE) enzyme GdpP (15). Bacterial strains that lack GdpP possess increased intracellular c-di-AMP levels, which leads to a growth defect but also to increased resistance to acid stress and β-lactam antibiotics (17,19-22). Previous attempts to construct a *dacA* deletion strain in *S. aureus* have failed, indicating that under standard laboratory conditions the production of c-di-AMP is important for the growth of *S. aureus* (23). However, *S. aureus* strains producing reduced levels of c-di-AMP have been described, which contain a point mutation in *dacA* resulting in a glycine to serine substitution at amino acid position 206 (*dacA_G206S_*)(20,21). These strains are viable, but have increased susceptibility to β-lactam antibiotics and acid stress (20,21). These results show that c-di-AMP is important for the growth of *S. aureus* under standard laboratory conditions.

Several c-di-AMP target proteins have been identified. In *S. aureus*, these include the proteins KtrA, PstA, KdpD, CpaA and OpuCA (24-29). KtrA is the regulatory gating component of the constitutively expressed Ktr potassium transport system and KdpD forms part of a two-component system required for the regulation of the Kdp transporter, a second potassium uptake system (25,30). PstA is a PII-like signal transduction protein, the cellular function of which is still unknown, even though several crystal structures are available (27,31,32). CpaA is a predicted cation/proton antiporter and OpuCA is the ATPase component of the carnitine ABC transporter OpuC (24,26,28). Interestingly, none of these uncovered target proteins were found to be essential for the growth of *S. aureus* under standard laboratory conditions (24-27,33). Thus, why c-di-AMP is important for the growth of *S. aureus*, and how the production and degradation of c-di-AMP is regulated, is yet to be answered.

Recently it was shown that *dacA* is essential for the growth of *Listeria monocytogenes* in rich medium, but dispensable when bacteria are grown in defined minimal medium (34). In the same study, it was reported that in the absence of c-di-AMP synthesis, the stringent response alarmones (p)ppGpp accumulate to toxic levels, preventing the growth of a *dacA* mutant. Consistent with this, *dacA* was dispensable in an *L. monocytogenes* strain lacking the three (p)ppGpp synthases RelA, RelP and RelQ (34) and hence unable to produce (p)ppGpp. More recently, it was shown that in the absence of c-di-AMP, the tricarboxylic acid (TCA) cycle intermediate citrate accumulates to toxic levels in *L. monocytogenes* (35). As TCA cycle activity is an important source for the generation of NADH (and therefore membrane potential) during aerobic respiration and is also intimately linked with amino acid metabolism (36), misregulation of TCA cycle activity, as observed in the absence of c-di-AMP in *L. monocytogenes*, will affect the function of many different cellular processes. In addition to mutations that prevent the production of (p)ppGpp, inactivating mutations that allow *L. monocytogenes* to grow in the absence of c-di-AMP were identified in the operons coding for the oligopeptide permease system Opp, the predicted glycine betaine transporter Gbu and the c-di-AMP target proteins CbpB and PstA (35). Based on additional work the authors suggested that increased uptake of osmolytes and peptides in the absence of c-di-AMP leads to an increase in the internal osmotic pressure and hence large differences between internal and external pressure (35). c-di-AMP has also been reported to be essential for growth of *Bacillus subtilis* (37,38). However, in contrast to *L. monocytogenes*, growth in chemically defined medium alone did not bypass the essentiality of c-di-AMP in *B. subtilis* (37). Only by reducing the concentration of potassium in the medium was the essentiality of c-di-AMP bypassed, suggesting that the lack of c-di-AMP leads to the extensive uptake of potassium and growth inhibition (37).

In the current study, we investigated the importance of c-di-AMP for the growth of *S. aureus*. We show that c-di-AMP production is dispensable for the growth of *S. aureus* in chemically defined medium and in rich medium supplemented with additional sodium or potassium chloride. In addition, we found that inactivating mutations in a glycine betaine uptake system and predicted amino acid transporter bypass the requirement of c-di-AMP for the growth of *S. aureus* in rich medium, further highlighting a key role of c-di-AMP in osmotic regulation. In addition, we found that mutations in genes encoding central aerobic respiration enzymes can bypass the requirement for c-di-AMP and that c-di-AMP is dispensable for the growth of *S. aureus* under anaerobic conditions. Taken together these results highlight an important function for c-di-AMP during aerobic but not anaerobic growth.

## RESULTS

### *S. aureus* strains with altered c-di-AMP levels show changes in membrane potential and cell size

Experimental evidence suggests that c-di-AMP negatively regulates potassium uptake in bacteria (24,37,39-41). Potassium is required for many cellular processes, including osmotic regulation and regulation of the membrane potential. Previous work has shown that an *S. aureus* strain lacking the constitutively expressed Ktr potassium uptake system has a hyper-polarized membrane (30). Hence, it is expected that c-di-AMP, through its control of potassium uptake, has an important role in regulating the membrane potential in bacteria. To test this hypothesis, we measured the membrane potential of the methicillin-resistant *S. aureus* (MRSA) USA300 strain LAC* (WT) and the isogenic mutant strains LAC\**gdpP::kan* (*gdpP*), which produces high levels of c-di-AMP, and LAC\**dacA_G206S_* (*dacA_G206S_*), which produces low levels of c-di-AMP, using the fluorescent membrane potential indicator dye 3, 3’-diethyloxacarbocyanine iodide [DiOC_2_(3)] and a FACS-based method. The green fluorescence emitted by this dye is dependent on cell size, while the red fluorescence is dependent on both cell size and membrane potential. Therefore, the ratio of red to green fluorescence gives a measure of the membrane potential that is largely independent of cell size (42). The *gdpP* mutant, which contains high levels of c-di-AMP and hence is expected to have lower cellular potassium levels, showed a statistically significant increase in membrane potential, while the low-level c-di-AMP strain *dacA_G206S_* showed a slightly, although not statistically significant, reduced membrane potential when compared to the WT strain (Fig. 1A and 1B). While performing the FACS experiments, apparent differences in the cell size between the strains were noted. This is in agreement with a previousley observed reduction in size of *gdpP* mutant cells (15). However, differences in cell size were also noted for the low c-di-AMP level producing strain *dacA_G206S_*. To investigate this in more detail, overnight cultures of WT, *gdpP* and *dacA_G206S_* strains were stained with a fluorescently-labelled vancomycin derivative that binds to peptidoglycan, thereby outlining the cell and allowing the size to be determined by fluorescence microscopy. The high-level c-di-AMP *gdpP* cells were smaller in size compared to the WT cells (Fig. 1C and 1D), in agreement with a previous report (15). Conversely, cells of the low-level c-di-AMP *dacA_G206S_* mutant strain were significantly larger (Fig. 1C and 1D). It is tempting to speculate that the larger cell size observed in a strain with reduced c-di-AMP levels is at least in part due to an increase in potassium and osmolyte uptake, leading to an increase in osmotic pressure at low c-di-AMP levels. This is consistent with additional findings presented below and previous speculations made by us and others (23,26,35,37,39,40).

**Figure 1.**
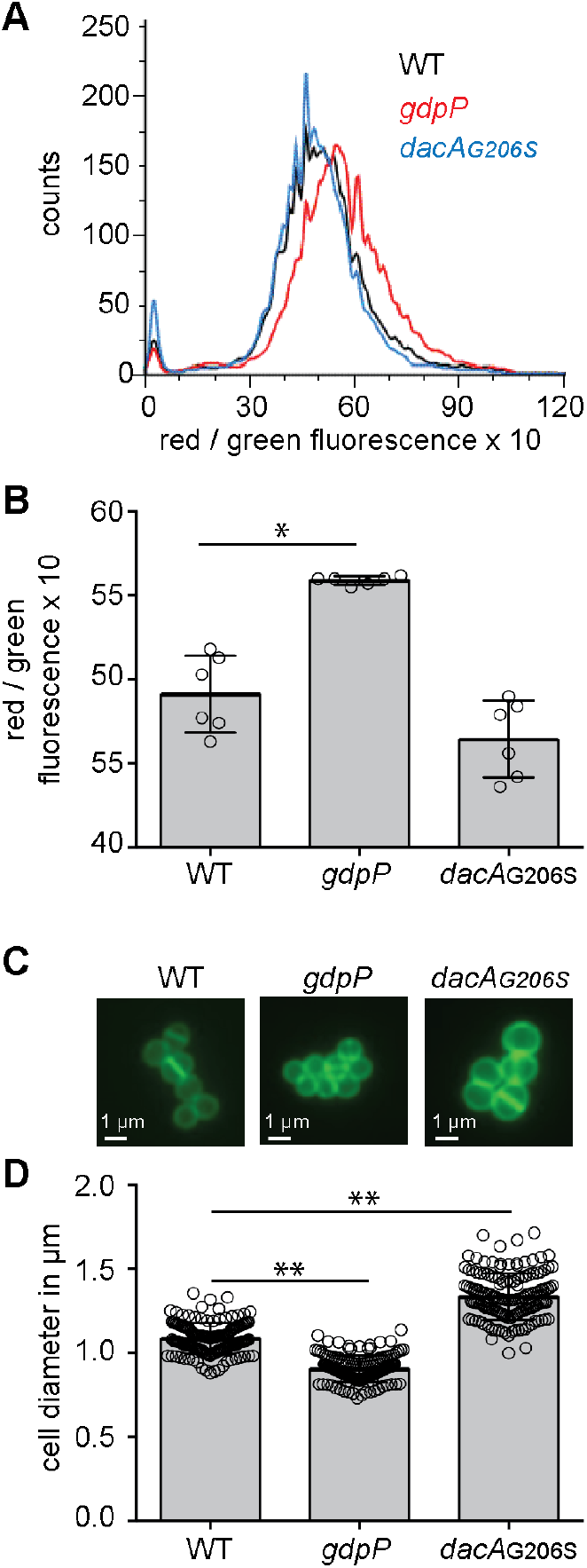
Variations in c-di-AMP levels impact the membrane potential and cell size of *S. aureus* cells. (A-B) Membrane potential measurement using a fluorescence-activated cell sorting (FACS)-based method. WT LAC* as well as the high c-di-AMP *gdpP* and low c-di-AMP *dacA_G2_o_6S_* mutant strains were grown overnight in TSB medium. Cells were washed and mixed with DiOC_2_(3) and the green and red fluorescence intensities detected using a FACSCalibur cytometer. The fluorescence intensities of 10000 gated events were recorded at the height of their emission peak. The ratio of red / green fluorescence x 10 was calculated for each event using the FlowJo V7 software and (A) representative histograms of cell counts versus fluorescence ratio are shown. (B) The mean values of red / green fluorescence x 10 was determined from the histograms in (A) and the averages and standard deviations (SDs) from six biological replicates plotted. (C-D) Bacterial cell size determination by microscopy. WT, *gdpP* and *dacA_G206S_* mutant strains were grown overnight in TSB medium. Culture aliquots were stained with Vancomycin-BODIPY and cells imaged using a fluorescence microscope. (C) Representative images of WT, *gdpP* and *dacA_G206S_* cells are shown. (D) The bacterial cell diameters were determined by drawing a line through the middle of nondividing cells using Image J. 150 cells were measured (3 biological replicates with 50 cells each) and the average cell diameters in μm and SDs were plotted. Statistical analysis was performed in Prism (GraphPad) using a Kruskal-Wallis test followed by a Dunn’s multiple comparison test. Adjusted p-values <0.05 are indicated by a single asterisk (*) and adjusted p-values < 0.01 by a double asterisk (**).

### DacA is dispensable for the growth of *S. aureus* in chemically defined medium

Previous attempts to delete the *dacA* gene in *S. aureus* under standard growth conditions in Tryptic Soy Broth (TSB) medium have been unsuccessful, indicating that c-di-AMP is important for the growth of *S. aureus* in rich medium (23). To determine if a *dacA* mutant could be obtained in *S. aureus* in chemically defined medium (CDM), a standard allelic exchange procedure was used in an attempt to replace the *dacA* gene with the *aph3* gene that confers kanamycin resistance, with the final steps performed in CDM. Using this approach, the strain LAC\**dacA::kan* (referred to as *dacA* mutant strain) was successfully constructed. To assess the growth characteristics of the *dacA* mutant strain, the plating efficiencies of this strain and the isogenic WT control strain on Tryptic Soy Agar (TSA) plates and on CDM plates were determined (Fig. 2A and 2B). This revealed that the *dacA* mutant strain had a severe growth defect on TSA plates but not on CDM plates; plating efficiencies were reduced nearly 5 logs on TSA plates as compared to CDM plates (Fig. 2B). Taken together these results show that *dacA* is important for the growth of *S. aureus* in TSB but dispensable for growth in CDM.

**Figure 2.**
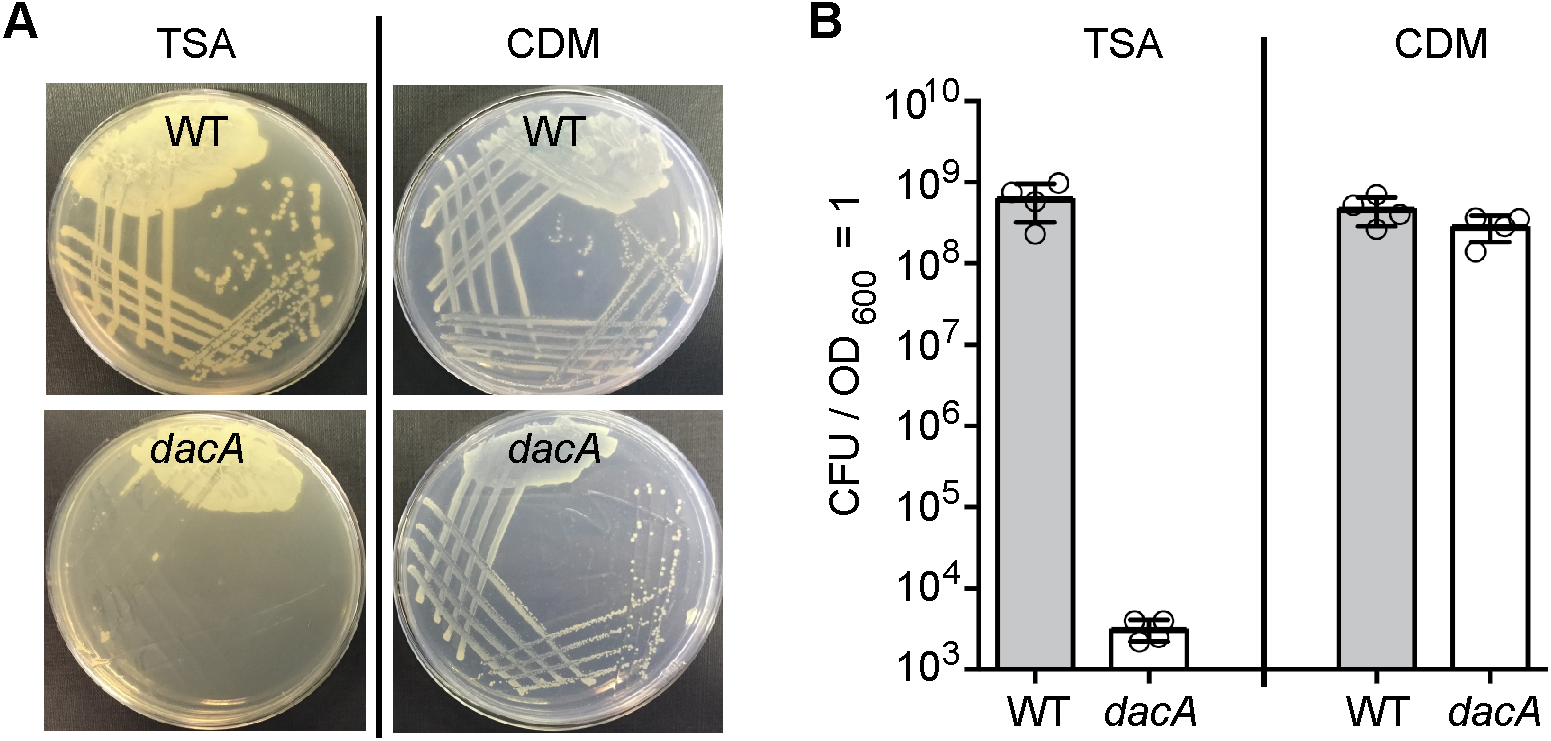
The *S. aureus dacA* mutant can grow in CDM but not in TSB medium. (A) Bacterial growth on agar plates. WT LAC* and the isogenic *dacA* mutant were streaked on TSA or CDM plates and images of plates taken after overnight incubation at 37°C. (B) Plating efficiencies. Bacterial suspensions were prepared for the WT and *dacA* mutant strains and appropriate dilutions spread on TSA or CDM plates and colony forming units (CFUs) per ml culture per OD_600_ unit determined and plotted. The average values and SDs from four biological samples were plotted.

### Mutations in genes coding for transporters and respiration-related genes suppress the growth defect of the *S. aureus dacA* mutant strain in rich medium

When higher cell density suspensions of a *S. aureus dacA* mutant culture were plated on TSA plates, a number of colonies were obtained. To investigate this further, several of these colonies referred to as suppressor strains LAC* *dacA::kan*-S1 to LAC\**dacA::kan*-S18 (or short S1 and S18) were picked and re-streaked on TSA plates to confirm their ability to grow in rich medium.The absence of c-di-AMP in cellular extracts was confirmed for several of these strains (S1, S2 and S4) using an LC-MS/MS approach (Fig. 3A). Next, a whole-genome sequencing strategy was employed on 17 suppressor strains (S1-S4 and S6-S18) to identify potential compensatory mutations. When compared to the genome sequence of WT LAC* (20), the *dacA* gene was deleted in all suppressor strains and one additional SNP at genome position 1622937, which is within the *xseA* gene, was observed in all suppressor strains, with the exception of strain S4. The mutation in this gene was presumably acquired during the strain construction process and strain S4 was likely derived from an earlier or different passage during the allelic exchange procedure (Table 1). In addition, one or two unique mutations were found in 14 of the 17 sequenced strains. Several strains had mutations at different locations within the same gene or within genes coding for proteins required for the same cellular processes, suggesting that mutations in these genes rather than the SNP in *xseA* are responsible for the growth rescue phenotype (Table 1). More specifically, mutations were identified in *opuD* (*SAUSA300_1245*) and *alsT* (*SAUSA300_1252*) coding for the predicted glycine-betaine transporter OpuD and the predicted alanine-proton symporter AlsT, respectively (Fig. 3B). In addition, several strains contained mutations in *hepS* (*SAUSA300_1359*), *ctaA* (*SAUSA300_1015*), *hemB* (*SAUSA300_1615*) or *qoxB* (*SAUSA300_0962*), coding for proteins required for the respiration process (Fig 3B, Table 1). A simplified view of the aerobic respiration chain in *S. aureus* and the function of HepS, CtaA, HemB and QoxB is presented in Fig. 3C and a more detailed description of their function will be provided in a later section.

**Figure 3.**
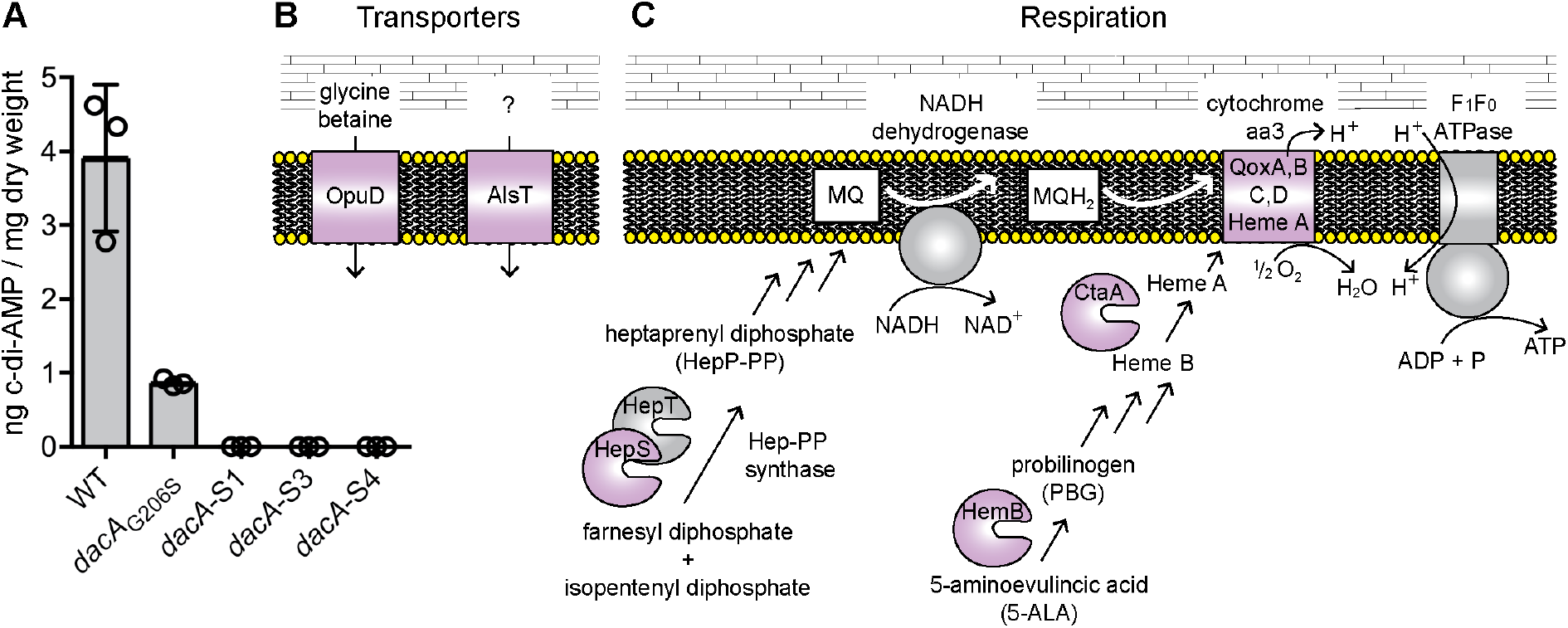
The *S. aureus dacA* suppressor strains do not produce c-di-AMP but acquire mutations in genes coding for transport or respiration related proteins. (A) Determination of cellular c-di-AMP levels by LC-MS/MS. Cell extracts (three biological replicates) were prepared from the suppressor strains dacA-S1, *dacA-S2* and *dacA-S4* and the production of c-di-AMP assessed by LC-MS/MS. The extracts were prepared and analyzed at the same time as WT LAC* and the low-level c-di-AMP *dacA_G206S_* strain as reported in a previous study (20). No c-di-AMP could be detected in extracts derived from strains *dacA-*S1, *dacA-S2* and *dacA-S4* and c-di-AMP levels determined for the WT and *dacA_G206S_* mutant as part of the previous study (20) are shown as controls. (B and C) Schematic representation of proteins whose genes were found to be mutated in *dacA* suppressor strains and their predicted cellular functions (B) Schematic representation and function of the OpuD and AlsT transporters. OpuD is a predicted (and as part of this study experimentally confirmed) glycine betaine and a reported weak proline transporter (43); AlsT is a predicted L-alanine/sodium symporter, but this substrate specificity could not be confirmed as part of this study. (C) Simplified view of the aerobic respiratory chain in *S. aureus*. The NADH dehydrogenase oxidizes NADH to NAD+ and electrons are transferred onto MQ to yield MQH_2_. The electrons are subsequently moved onto the heme A component of the cytochrome aa3 complex, composed of the QoxA, B, C, D components. Upon reduction of ½O_2_ to H_2_O by the cytochrome complex, protons (H+) are extruded and utilized by the F_1_F_0_-ATPase for the production of ATP. Suppressor mutations were identified in genes coding for HepS, HemB, CtaA (a membrane protein but shown for simplicity as soluble proteins) and QoxB, required for MQ, HemA and Cytrochome aa3 complex formation, respectively. The proteins for which gene variations were found in suppressor strains are indicated in purple in the schematics.

**Table 1:**
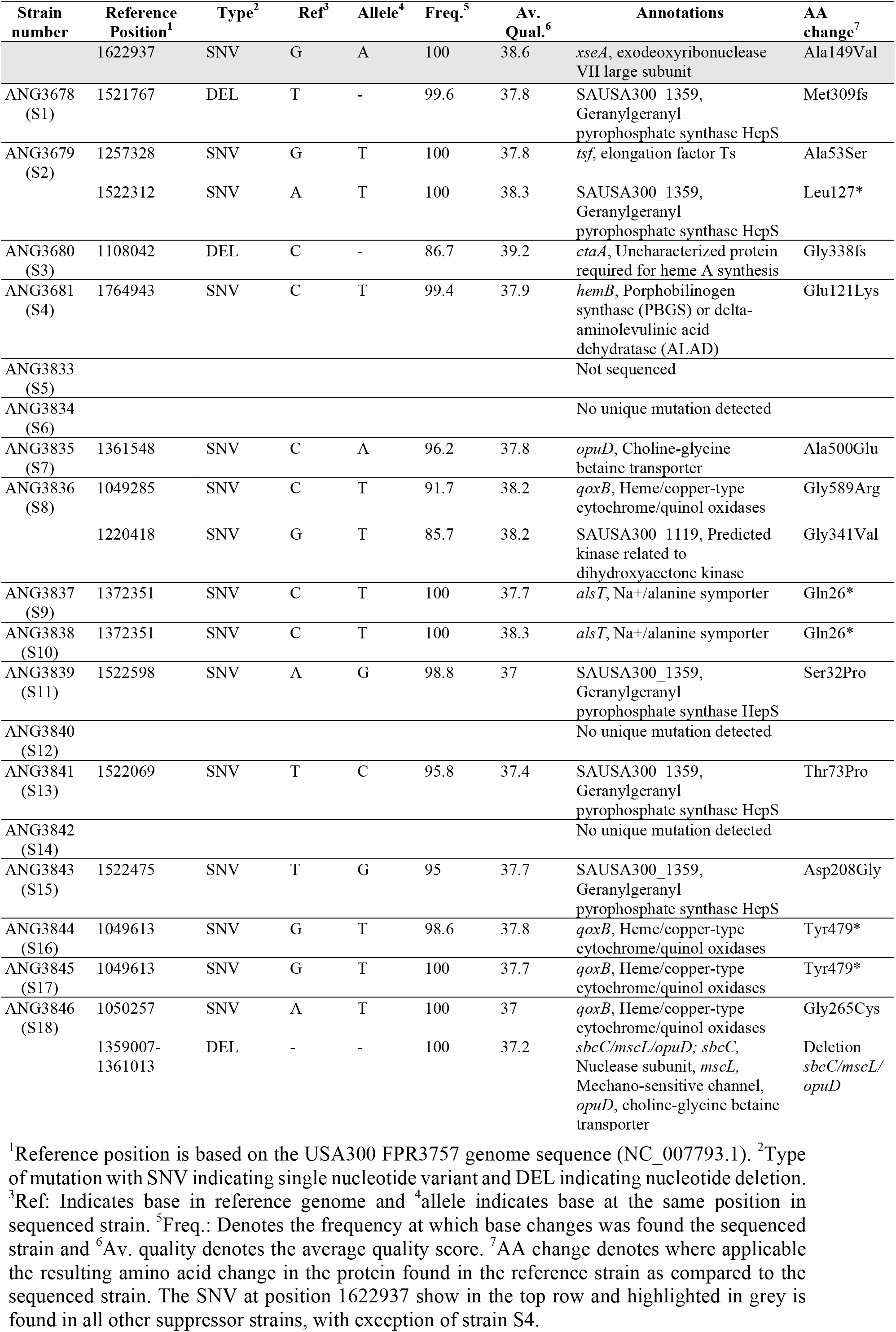
Sequence alterations in LAC\**dacA::kan* suppressor strains compared to LAC*

| Strain number | Reference Position<sup>1</sup> | Type<sup>2</sup> | Ref<sup>3</sup> | Allele<sup>4</sup> | Freq.<sup>5</sup> | Av. Qual.<sup>6</sup> | Annotations | AA change<sup>7</sup> |
| --- | --- | --- | --- | --- | --- | --- | --- | --- |
|  | 1622937 | SNV | G | A | 100 | 38.6 | *xseA*, exodeoxyribonuclease VII large subunit | Ala149Val |
| ANG3678 (S1) | 1521767 | DEL | T | - | 99.6 | 37.8 | SAUSA300_1359, Geranylgeranyl pyrophosphate synthase HepS | Met309fs |
| ANG3679 (S2) | 1257328 | SNV | G | T | 100 | 37.8 | *tsf*, elongation factor Ts | Ala53Ser |
|  | 1522312 | SNV | A | T | 100 | 38.3 | SAUSA300_1359, Geranylgeranyl pyrophosphate synthase HepS | Leu127* |
| ANG3680 (S3) | 1108042 | DEL | C | - | 86.7 | 39.2 | *ctaA*, Uncharacterized protein required for heme A synthesis | Gly338fs |
| ANG3681 (S4) | 1764943 | SNV | C | T | 99.4 | 37.9 | *hemB*, Porphobilinogen synthase (PBGS) or delta-aminolevulinic acid dehydratase (ALAD) | Glu121Lys |
| ANG3833 (S5) |  |  |  |  |  |  | Not sequenced |  |
| ANG3834 (S6) |  |  |  |  |  |  | No unique mutation detected |  |
| ANG3835 (S7) | 1361548 | SNV | C | A | 96.2 | 37.8 | *opuD*, Choline-glycine betaine transporter | Ala500Glu |
| ANG3836 (S8) | 1049285 | SNV | C | T | 91.7 | 38.2 | *qoxB*, Heme/copper-type cytochrome/quinol oxidases | Gly589Arg |
|  | 1220418 | SNV | G | T | 85.7 | 38.2 | SAUSA300_1119, Predicted kinase related to dihydroxyacetone kinase | Gly341Val |
| ANG3837 (S9) | 1372351 | SNV | C | T | 100 | 37.7 | *alsT*, Na<sup>+</sup>/alanine symporter | Gln26* |
| ANG3838 (S10) | 1372351 | SNV | C | T | 100 | 38.3 | *alsT*, Na<sup>+</sup>/alanine symporter | Gln26* |
| ANG3839 (S11) | 1522598 | SNV | A | G | 98.8 | 37 | SAUSA300_1359, Geranylgeranyl pyrophosphate synthase HepS | Ser32Pro |
| ANG3840 (S12) |  |  |  |  |  |  | No unique mutation detected |  |
| ANG3841 (S13) | 1522069 | SNV | T | C | 95.8 | 37.4 | SAUSA300_1359, Geranylgeranyl pyrophosphate synthase HepS | Thr73Pro |
| ANG3842 (S14) |  |  |  |  |  |  | No unique mutation detected |  |
| ANG3843 (S15) | 1522475 | SNV | T | G | 95 | 37.7 | SAUSA300_1359, Geranylgeranyl pyrophosphate synthase HepS | Asp208Gly |
| ANG3844 (S16) | 1049613 | SNV | G | T | 98.6 | 37.8 | *qoxB*, Heme/copper-type cytochrome/quinol oxidases | Tyr479* |
| ANG3845 (S17) | 1049613 | SNV | G | T | 100 | 37.7 | *qoxB*, Heme/copper-type cytochrome/quinol oxidases | Tyr479* |
| ANG3846 (S18) | 1050257 | SNV | A | T | 100 | 37 | *qoxB*, Heme/copper-type cytochrome/quinol oxidases | Gly265Cys |
|  | 1359007-1361013 | DEL | - | - | 100 | 37.2 | *sbcC/mscL/opuD*; *sbcC*, Nuclease subunit, *mscL*, Mechano-sensitive channel, *opuD*, choline-glycine betaine transporter | Deletion *sbcC/mscL/opuD* |

<sup>1</sup>Reference position is based on the USA300 FPR3757 genome sequence (NC\_007793.1). <sup>2</sup>Type of mutation with SNV indicating single nucleotide variant and DEL indicating nucleotide deletion.
<sup>3</sup>Ref: Indicates base in reference genome and <sup>4</sup>allele indicates base at the same position in sequenced strain. <sup>5</sup>Freq.: Denotes the frequency at which base changes was found the sequenced strain and <sup>6</sup>Av. quality denotes the average quality score. <sup>7</sup>AA change denotes where applicable the resulting amino acid change in the protein found in the reference strain as compared to the sequenced strain. The SNV at position 1622937 show in the top row and highlighted in grey is found in all other suppressor strains, with exception of strain S4.

### Growth, complementation and uptake analysis of *dacA* suppressor strains with mutations in *opuD* and *alsT*

The mutations obtained in *opuD* and *alsT* point to a role for c-di-AMP in regulating the osmolyte and amino acid/peptide concentration in *S. aureus*. Using suppressor strains S10 (*alsT*), S7 (*opuD*) and S18 (*qoxB*/Δ*opuD*), our further studies initially focused on the cellular function of the *S. aureus* OpuD and AlsT transporters. The *opuD* mutation in strain S7 (*opuD*) and the deletion of the complete *opuD* gene in strain S18 (*qoxB*/Δ*opuD*) indicates that inactivation of OpuD could lead to the observed growth rescue in rich medium. Of note, strain S18 also has a mutation in the *qoxB* gene, the function of which will be discussed later. Suppressor strains S9 and S10 (*alsT*) contained the same SNP in *alsT* that creates a stop codon truncating the encoded protein after 26 amino acids, indicating that inactivation of AlsT leads to the observed suppressor phenotype. Initially the growth of suppressor strains S10 (*alsT*), S7 (*opuD*) and S18 (*qoxB*/Δ*opuD*) as well as the WT LAC* and the original *dacA* mutant control strains in CDM and TSB medium was assessed in detail. As expected, all strains were able to grow in CDM (Fig. 4A). The *dacA* mutant showed the expected growth defect in TSB medium, but all three suppressor strains showed improved growth in TSB, confirming that these strains are *bona fide* suppressor strains (Fig. 4B). Thus, it is likely that loss of either AlsT or OpuD activity leads to the observed rescue of growth of the *dacA* mutant in rich medium. Hence, expression of a wild-type copy of *alsT* or *opuD* in the respective suppressor strains should lead to a growth inhibition in TSB. While all our attempts to introduce a plasmid for the expression of *alsT* in the respective suppressor strains failed, plasmid piTET-opuD was successfully introduced into suppressor strains S7 (*opuD*) and S18 (*qoxB/ΔopuD)*. Plasmids piTET-*opuD* integrates into the *geh* locus of *S. aureus* and allows for anhydroteracycline (Atet) inducible *opuD* expression. As a further control, the empty vector piTET was also introduced into WT LAC* and the suppressor strains. Successful complementation should result in a reduction of the plating efficiency of strains S7 (*opuD*) and S10 (*qoxB/ΔopuD*) containing plasmid piTET-opuD when plated on TSA plates supplemented with 200 ng/ml Atet. This was indeed the case; plating efficiencies of the complemented strains were reduced by 2-3 logs as compared to the WT or empty vector control strains (Fig. 4C). The plating efficiencies were not reduced to the same levels of the original *dacA* mutant strain, indicating only partial complementation, potentially owing to insufficient levels of *opuD* expression (Fig. 4C). Taken together, this growth analysis indicates that the *dacA* suppressor strains containing inactivating mutations in *opuD* and *alsT* show improved growth in rich medium, with complementation analysis providing particularly strong evidence in the case of the *opuD* mutant strains.

**Figure 4.**
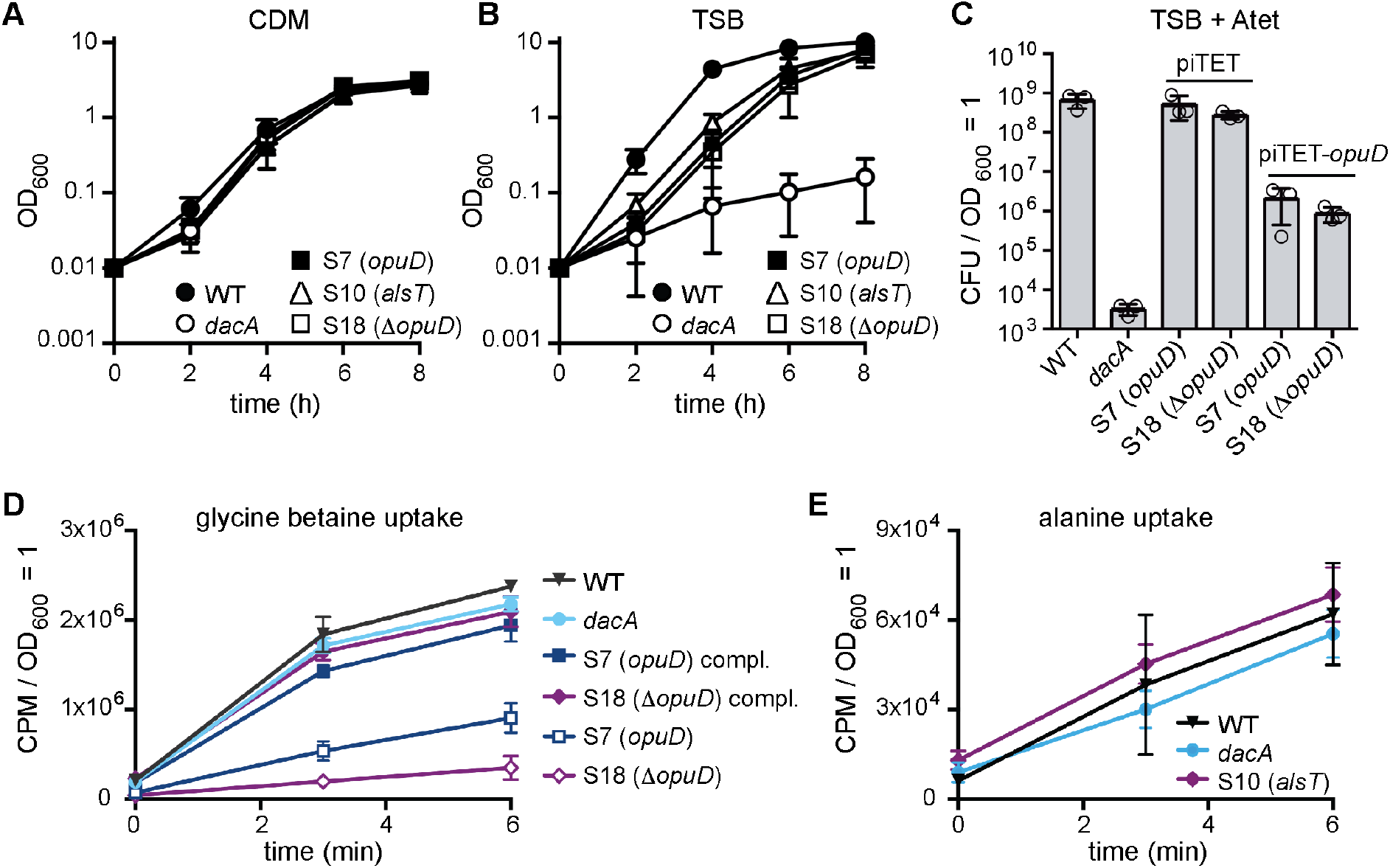
*S. aureus dacA* suppressors with mutations in *opuD and alsT* have improved growth in TSB. (A-B) Bacterial growth curves. WT LAC*, the *dacA* mutant and suppressor strains S7 (*opuD*), S10 (*alsT*), and S18 (*qoxB/*Δ*opuD*) [or short S18 (Δ*opuD)]* were propagated in (A) CDM or (B) TSB medium and their growth monitored by taking OD_600_ readings. The average values and SDs from three independent experiments were plotted. (C) Genetic complementation of *opuD* mutants. Bacterial suspensions were prepared for WT, the *dacA* mutant as well as suppressor strains S7 (*opuD*) and S18 (*qoxB/*Δ*opuD*) [or short S18 (Δ*opuD)]* containing the empty piTET vector or the complementation plasmid piTET-opuD. Appropriate culture dilutions were plated on TSA plates containing 200 ng/ml Atet and CFUs per ml culture per OD_600_ unit determined and plotted. The average values and SDs from three experiments were plotted. (D) *opuD* mutants are defective in glycine betaine (GB) uptake. WT, the *dacA* mutant and the suppressor strains S7 (*opuD*) and S18 (*qoxB/*Δ*opuD*) [or short S18 (Δ*opuD)]* containing the empty piTET vector or the plasmid piTET-opuD were grown to mid-log phase in CDM supplemented with 200 ng/ml Atet. For uptake assays, radiolabeled GB was added to culture aliquots, samples removed and filtered at the indicated time points and the radioactivity accumulated in the cells measured. The average values and SDs from four experiments were plotted. (E) Alanine uptake assays. WT, *dacA* and S10 (*alsT*) strains were grown to midlog phase in CDM containing half the L-alanine concentration as in the standard medium. Bacterial suspensions were prepared and radiolabeled alanine added to the cultures. Sample aliquots were removed at the indicated time points, filtered and the radioactivity accumulated in the cells measured. The average values and SDs from four experiments were plotted.

OpuD is annotated as a glycine betaine transporter belonging to the Betaine-Carnitine-Choline Transporter (BCCT) family whereas AlsT is annotated as an amino acid carrier protein and often is more specifically referred to as an alanine/sodium symporter. However, to the best of our knowledge neither OpuD nor AlsT have been tested as glycine betaine or alanine transporters in *S. aureus*. In a recent study, OpuD has been implicated as a weak and low affinity proline uptake system in *S. aureus* (43). To determine potential substrates for the *S. aureus* OpuD and AlsT transporters, uptake assays were performed with the WT, *dacA* mutant and the *opuD* and *alsT* suppressor strains using either radiolabeled glycine betaine or alanine as substrate. Glycine betaine uptake was severely reduced in suppressor strains S7 (*opuD*) and S18 (*qoxB*/Δ*opuD*) compared to the WT strain (Fig. 4D). Glycine betaine uptake was restored to almost WT levels in the complemented *opuD* strains (Fig. 4D). Hence, OpuD appears to function as main glycine betaine transporter in *S. aureus* under our growth conditions. In contrast, no difference in alanine uptake was detected between strain S10 (*alsT*), which contains a stop codon in AlsT, the WT and *dacA* mutant strains (Fig. 4E). This indicates that AlsT is unlikely the main alanine uptake system in *S. aureus* under our assay conditions.

### Glycine betaine and peptides have a negative impact whereas salt has a positive impact on the growth of the *S. aureus dacA* mutant

While we have not been able to determine the substrate specificity of AlsT, based on its sequence and homology to other transporters it seems likely that this protein is involved in the uptake of an amino acid or peptides. Therefore, the data presented thus far indicate that uptake of glycine betaine by OpuD and perhaps some amino acid or peptide by AlsT contribute to the observed growth defect of the *dacA* mutant in rich medium. Consequently, the addition of glycine betaine and tryptone as extra peptides source to CDM might have a negative impact on the growth of the *dacA* mutant. To test this, the growth of WT LAC*, the *dacA* mutant and the S7 (*opuD*) and S10 (*alsT*) suppressors in CDM and in CDM supplemented with 1 mM glycine betaine or 1% tryptone was compared. The *dacA* mutant and suppressor strains grew with the same growth kinetics in CDM (Fig. 5A), but supplementing the CDM with glycine betaine (Fig. 5B) or tryptone (Fig. 5C) reduced the growth of only the *dacA* mutant; the growth of the suppressor strains was unaffected. It is of interest to note that the *opuD* and *alsT* suppressor strains were insensitive to the addition of both, glycine betaine and tryptone, suggesting that osmolyte and peptide uptake converge at one point to impede the growth of the *dacA* mutant. Glycine betaine and several different amino acids are known intracellular osmolytes. For *L. monocytogenes*, it has been proposed that a high intracellular osmolyte concentration in the *dacA* mutant strain leads to an increased internal osmotic pressure and that the resulting imbalance between the internal and external pressure results in the observed growth inhibition. Consistent with this model, increasing the external osmotic pressure through the addition of NaCl or KCl to the growth medium rescued the growth of an *L. monocytogenes dacA* mutant. To test if this is also the case in *S. aureus*, we plated the WT and *dacA* mutant strains on TSA plates containing increasing NaCl or KCl concentrations. Indeed, increasing the osmolarity of the medium by the addition of NaCl or KCl rescued the growth of the *dacA* mutant strain with greatest improvement of growth seen in TSB supplemented with 0.8 M NaCl or 0.8 M KCl (Fig. 5D and 5E). However, at very high NaCl or KCl concentrations of 2 M, the *dacA* mutant demonstrated again a significant plating defect (Fig. 5D and 5E). Compared to other bacteria, *S. aureus* is extremely tolerant to osmotic stress and as shown here can grow on agar plates containing 2 M NaCl. For *S. aureus* to be able to divide under these conditions, specific osmotic stress adaption processes are activated and as previously reported a large number of transcriptional changes occur (44,45). While intermediate salt concentrations can rescue the growth of a *dacA* mutant, the strain might not be able to mount the appropriate stress adaption response under extremely high osmolarity conditions, leading to the observed drop in plating efficiency. Taken together, these observations are in agreement with findings in *L. monocytogenes* (35) and point towards a common mechanism for the growth requirement of c-di-AMP in both organisms, where the bacteria are unable to maintain balanced intracellular osmolyte and amino acid concentrations in the cell in the absence of c-di-AMP. On the other hand, in *B. subtilis* the accumulation of potassium leads in the absence of c-di-AMP to the growth defect (37).

**Figure 5.**
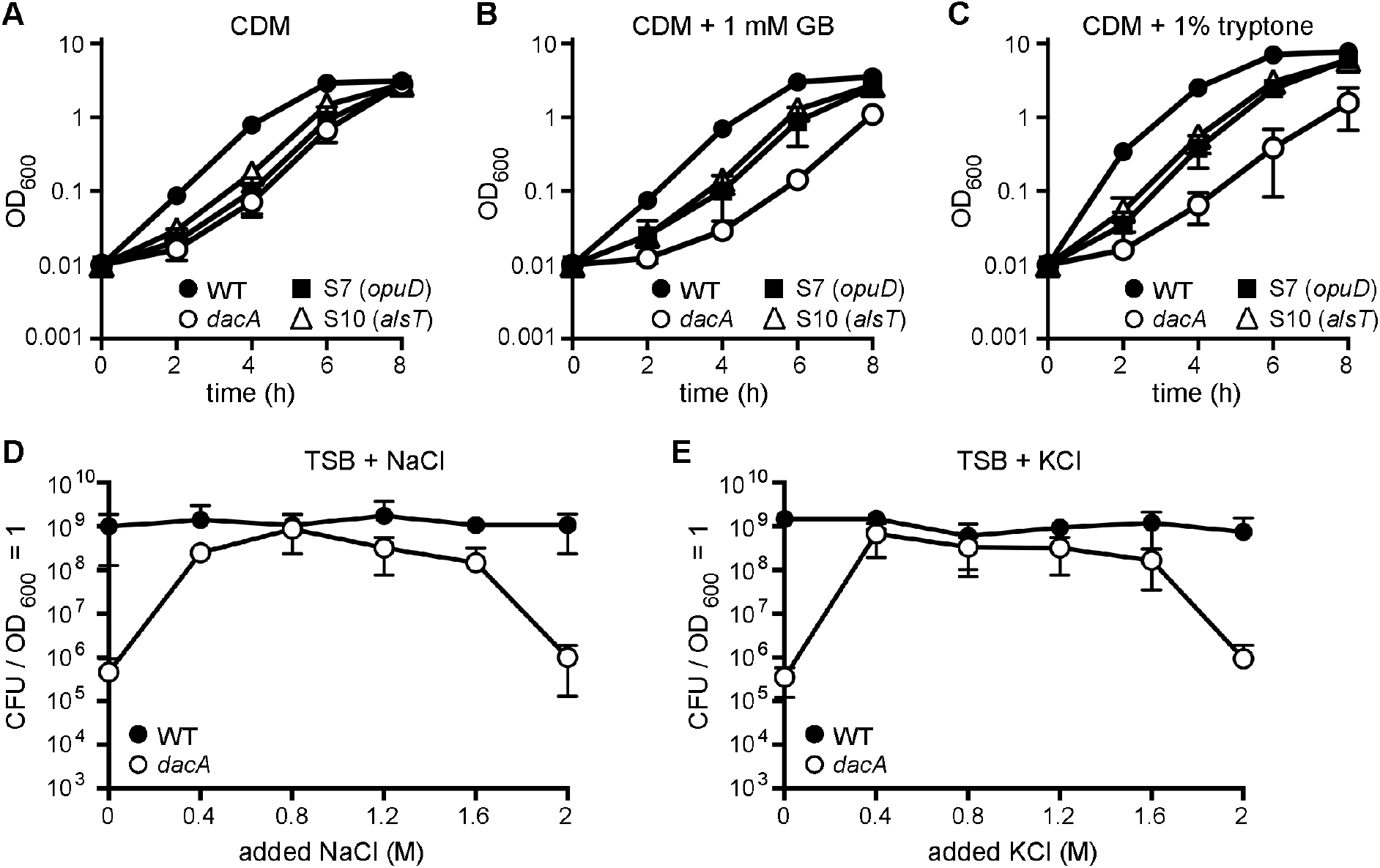
Glycine betaine and peptides inhibit, while salt improves the growth of the *dacA* mutant. (A-C) Bacterial growth curves. WT LAC*, the *dacA* mutant and S7 (*opuD*) and dacA-S10 (*alsT*) suppressor strains were propagated in (A) CDM, (B) CDM with 1 mM glycine betaine (GB) or (C) CDM with 1% tryptone and their growth monitored by determining OD_600_ readings. The average values and SDs from three independent experiments were plotted. (D-E) Plating efficiencies of *S. aureus* in TSB medium supplemented with increasing amounts of NaCl or KCl. Bacterial suspensions were prepared for WT LAC* and the *dacA* mutant strain and serial dilutions spotted on TSA or TSA plates containing the indicated concentrations of (D) NaCl or (E) KCl. The average CFUs per ml culture per OD_600_ unit and the SDs from three experiments were plotted.

### *S. aureus dacA* suppressor strains with mutations in genes coding for respiration-related proteins have improved growth in TSB medium that can be complemented chemically or genetically

While suppressor mutations in genes coding for transporters are similar to mutations previously observed in *L. monocytogenes*, the majority of *S. aureus dacA* suppressor strains contained mutations in genes encoding for proteins not previously associated with the c-di-AMP signaling pathways. Specifically, mutations were observed in *ctaA, hemB, hepS* and *qoxB*, which code for proteins required for respiration (Table 1 and Fig. 3C). When *S. aureus* respires aerobically, electrons are passed by a dehydrogenase (i.e. NADH dehydrogenase) onto menaquinones (MQ) to yield MQH_2_ (Fig. 3C). Next, the electrons are shuttled to the heme group within the cytochrome aa3 complex and subsequently used for the reduction of O_2_ to H_2_O. Protons (H^+^) are extruded during this process and the generated proton motive force (PMF) is used, amongst others, by the F_1_F_0_ ATPase to generate ATP. The predicted functions of HepS, HemB, CtaA and QoxB are as follows: HepS, together with HepT, likely forms a heptaprenyl-diphosphate synthase (HepP-PP synthase) and is responsible for the production of a precursor for MQ synthesis. HemB and CtaA are required for Heme A biosynthesis, which is the heme co-factor within the cytochrome aa3 complex. Lastly, QoxB is one of the protein components within the cytochrome aa3 complex, which represents one of the two main terminal oxidases used in *S. aureus* for respiration. Some of the mutations identified in *hepS, ctaA, hemB* and *qoxB* are predicted to result in either amino acid changes within important functional domains or introduce premature stop codons (Table 1), in both cases likely leading to inactive proteins. Hence aerobic respiration might be inhibited or reduced in these suppressor strains. Representative suppressor strains, namely S3 (ctaA), S4 (*hemB*), S13 (*hepS*) and S16 (*qoxB*) were chosen for further growth and subsequent complementation analysis. The mutations in *ctaA, hemB, hepS* or *qoxB* identified by the whole genome sequence approach were confirmed for these strains by fluorescence automated sequencing. Next, their growth was assessed and compared to that of WT LAC* and the original *dacA* mutant strain in TSB and CDM. Strains S3 (ctaA), S4 (*hemB*) and S16 (*qoxB*) grew as expected for *bona fide* suppressor strains, as these strains grew in both CDM and TSB medium (Fig. 6A and 6B). Their growth rate in TSB was somewhat reduced compared to the WT control strain, however, the growth was significantly improved when compared to the *dacA* mutant (Fig. 6A and 6B). While the suppressor strain S13 (*hepS*) was able to grow on CDM and TSA plates, it grew very poorly in either medium in liquid culture (Fig. 6A and 6B). This indicates that *hepS* mutations may only be able to bypass the *dacA* growth requirement on solid but not in liquid medium and therefore this strain was not included in any further analysis. To confirm that inactivation of the proteins encoded by *ctaA, hemB* and *qoxB* leads to the suppression of the *dacA* growth phenotype, we performed complementation analyses where successful complementation should result in a plating defect on TSA plates. A *hemB* mutant is unable to produce heme and this defect can be complemented chemically by addition of hemin to the medium, which bypasses the requirement for HemB (46,47). While the plating efficiency of strain S4 (*hemB*) on TSA was similar to that of the WT, in the presence of 10 μM hemin in the plates the plating efficiency decreased drastically (Fig. 6C). A genetic complementation strategy was used for suppressor strains S3 (*ctaA*) and S16 (*qoxB*) as well as strain S18 (*qoxB/opuD*), which contained both a mutation in *qoxB* and a deletion of the *opuD* region. To this end plasmids piTET-ctaA and piTET-qoxB were constructed and introduced into the respective suppressor strains. Either complete or partial complementation was observed upon expression of *ctaA* or *qoxB*, as determined by plating efficiencies on TSA plates containing 200 ng/ml Atet (Fig. 6D and 6F). Taken together this complementation analysis strongly suggests that inactivation or reduced activity of *ctaA, hemB* or *qoxB* results in the observed suppression of the *dacA* growth requirement under aerobic conditions in rich medium.

**Figure 6.**
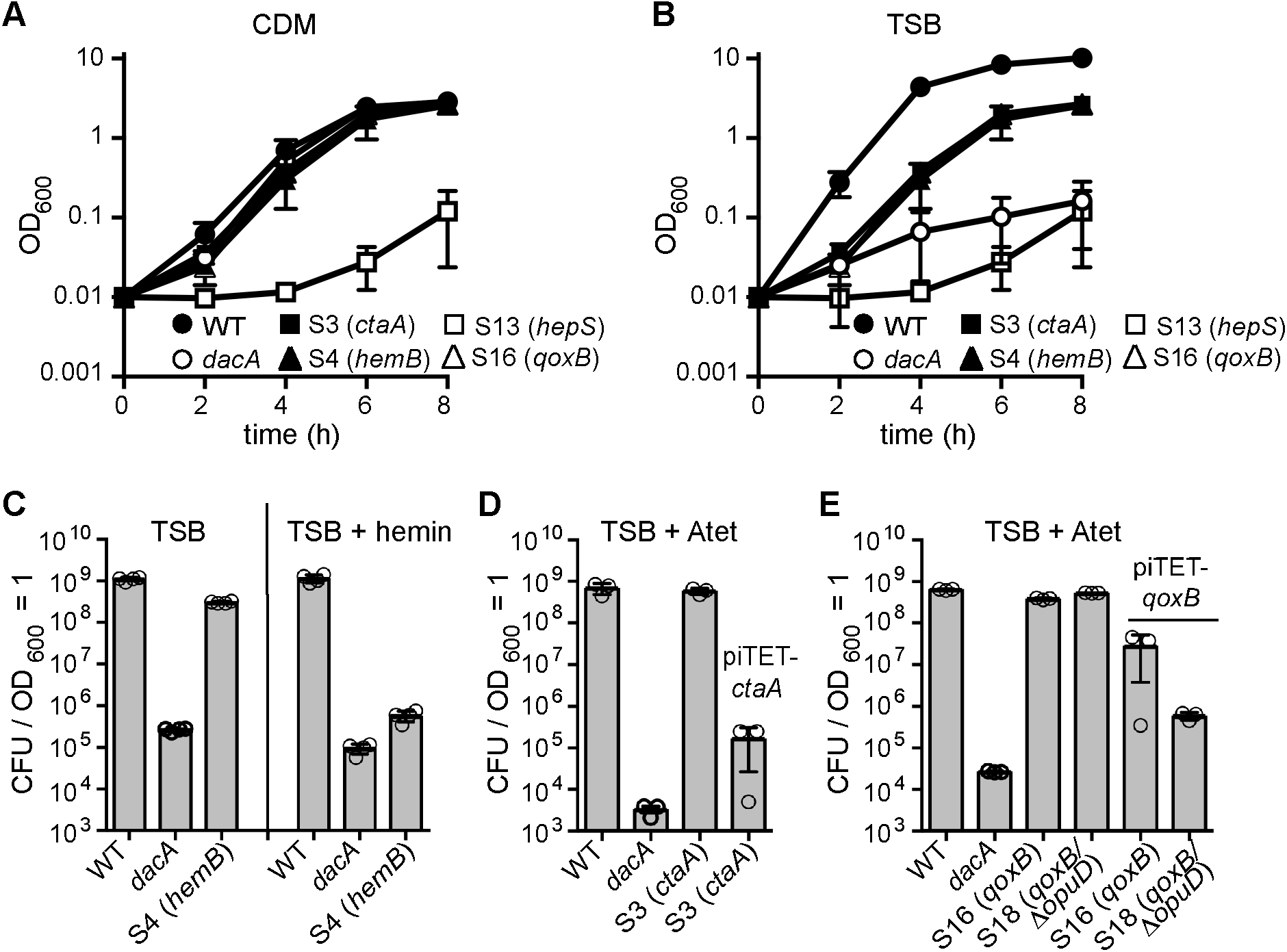
*S. aureus dacA* suppressors with mutations in *ctaA, hemB* or *qoxB* have improved growth in TSB that can be complemented chemically or genetically. (A-B) Bacterial growth curves. WT LAC*, the *dacA* mutant and the suppressor strains S3 (ctaA), S4 (*hemB*), S13 (*hepS*) and S16 (*qoxB*) were propagated in (A) CDM or (B) TSB medium and their growth monitored by OD_600_ readings. The average values and standard deviations of three independent experiments were plotted. (C) Chemical complementation analysis of the *hemB* mutant. Bacterial suspensions were prepared for WT LAC*, the *dacA* mutant and suppressor strain S4 (*hemB*) and appropriate dilutions plated on TSA or TSA plates containing 10 μM hemin. CFUs per ml culture per OD_600_ unit were determined and plotted. (D-E) Genetic complementation analysis of (D) *ctaA* and (E) *qoxB* mutants. Bacterial suspensions were prepared for WT LAC*, the *dacA* mutant as well as (C) suppressor strain S3 (ctaA) with or without the complementation plasmid piTET-ctaA or (D) suppressor strains S16 (*qoxB*) and S18 (*qoxB/*Δ*opuD*) with or without the complementation plasmid piTET-qoxB. Dilutions were plated on TSA plates containing 200 ng/ml Atet and the average CFUs per ml culture per OD_600_ unit and SDs from three experiments plotted.

### Utilization of the main terminal oxidase system Qox is decreased in the *S. aureus dacA* suppressor strains

*S. aureus* posesses two terminal oxidases, the Qox system (also referred to as cytochrome aa3) and the Cyd system (also referred to as cytochrome bd) (Fig. 7A). The presence of mutations in respiration-related genes indicated that alterations in bacterial aerobic respiration might impact the requirement for c-di-AMP for bacterial growth. To assess bacterial respiration, the oxygen consumption rates were determined for the WT *S. aureus* strain and the suppressor strains S3 (*ctaA*), S4 (*hemB*) and S16 (*qoxB*) using a Clark-type oxygen electrode. As controls, the oxygen consumption rates were also measured for *S. aureus* strains with transposon insertions in either *ctaA* or *qoxB*, which lack a functional Qox system, and a *hemB* deletion strain (Δ*hemB*), which is unable to produce functional cytochrome aa3 and cytochrome bd and hence lacks both a functional Qox and a functional Cyd system (Fig. 7A). The control strains with transposon insertions in *ctaA* and *qoxB* displayed an aerobic respiration rate similar to that of the WT strain (Fig. 7B), whereas the *hemB* deletion strain showed the expected reduction in oxygen consumption (Fig. 7B). These findings are consistent with previously published results, showing that only simultaneous inactivation of both terminal oxidases leads to a reduced membrane potential in *S. aureus* (10). The *dacA* suppressor strains S3, S4 and S16, with mutations in *ctaA, hemB* and *qoxB*, respectively, showed slightly reduced aerobic respiration rates as compared to the WT strain, but were still able to consume oxygen (Fig. 7B). The reduction in the oxygen consumption rate of suppressor strain S4 (*hemB*) is not as drastic as that of the *ΔhemB* control strain, suggesting that the point mutation in *hemB* in S4 strain only partially impairs the enzymatic activity of HemB. As suppressor strains S3 and S16 contained mutations leading to premature stop codons in CtaA and QoxB, respectively, it seemed likely that the main terminal oxidase (the Qox system) is defective in these suppressor strains and that these strains respire using only the Cyd system. To test this experimentally, a *cydA* transposon mutation (*cydA::tn*) was introduced into suppressor strains S3 (ctaA) and S16 (*qoxB*) and the oxygen consumption rates of the resulting strains compared to that of the WT and a *cydA::tn* mutant, which lacks only the Cyd system. As expected, the WT and *cydA::tn* mutant displayed similar respiration rates (Fig. 7C). However, the *dacA* suppressor strains S3 (*ctaA*) also containing the *cydA::tn* mutation displayed a drastically reduced oxygen consumption rate and strain S16 (*qoxB*) *cydA::tn* was basically unable to consume oxygen (Fig. 7C). Taken together, these data show that while the *dacA* suppressor strains with mutations in respiration-related genes can still consume oxygen, respiration in these strains is either reduced or proceeds primarily the Cyd terminal oxidase and rather than the proton-pumping Qox terminal oxidase system.

**Figure 7.**
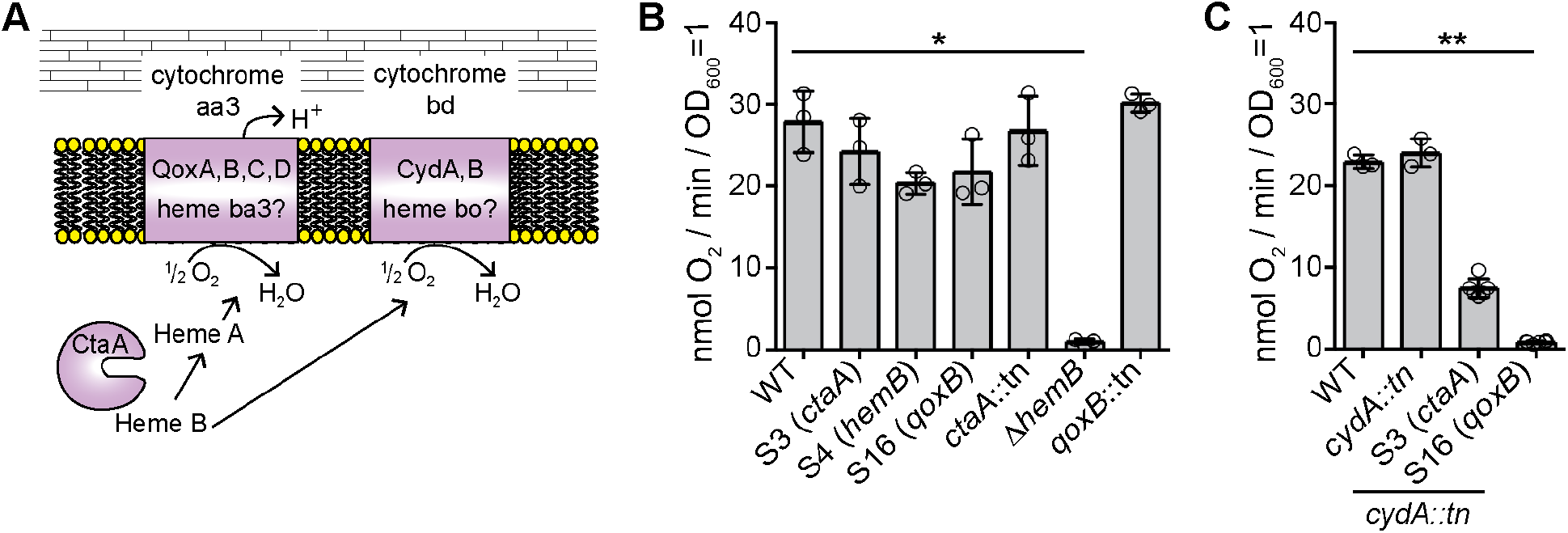
Oxygen consumption rates of WT, mutant and *dacA* suppressor strains. (A) Schematic representation of the two *S. aureus* terminal oxidases. The main terminal oxidase Qox (also referred to as cytochrome aa3) is composed of the proteins QoxA-QoxD and has been suggested to contain Heme A and Heme B as cofactors (10) and requires the CtaA protein (a membrane protein but shown for simplicity as soluble protein) for Heme A biosynthesis. The second terminal oxidase Cyd (also referred to as cytochrome bd) is composed of the proteins CydA and CydB and has been suggested to contain Heme B and Heme O (10) and therefore does not require CtaA for its synthesis. (B-C) Oxygen consumption rates of WT, mutant and *dacA* suppressor strains. The indicated *S. aureus* strains were grown to mid-log phase, washed and the oxygen consumption rates of concentrated culture aliquots determined following the addition of glucose. The oxygen consumption rates are plotted as nmol O_2_ consumed per min per OD_600_ = 1. For each strain, 3 to 6 biological replicates were used and the average values and standard deviations of the oxygen consumption rates were plotted. Statistical analysis was performed in Prism (GraphPad) using a Kruskal-Wallis test followed by a Dunn’s multiple comparison test. Adjusted p-values <0.05 are indicated by a single asterisk (*) and adjusted p-values < 0.01 by a double asterisk (**).

### *S. aureus* strains with altered c-di-AMP levels show changes in endogenous reactive oxygen species production

During aerobic respiration, endogenous reactive oxygen species (ROS) are produced that at high levels cause DNA damage and also result in lipid and protein peroxidation (48). To test if differences in c-di-AMP levels impact endogenous ROS production, its production in the WT, the high-level c-di-AMP *gdpP* and the low-level c-di-AMP *dacA_G206S_* mutants was assessed using the ROS sensitive fluorescence indicator dye 2’,7’-dichlorofluorescein diacetate and a previously published method (49,50). Of note, as the *dacA* mutant cannot grow in TSB medium under aerobic conditions, only the low c-di-AMP level *dacA_G206S_* mutant stain was included in this analysis. Also, the obtained fluorescence values were normalized based on the number of colony forming units (CFUs) rather than OD600 readings, as the WT, *gdpP* and *dacA_G206S_* mutant strains differ in cell size (see Figs. 1C and 1D), which impacts OD600 readings. As controls, *S. aureus* strains SH1000 and the isogenic catalase (KatA) and alkyl hydroperoxide reductase (AhpC) mutant SH1000Δ*katA*Δ*ahpC* were used. As expected, strain SH1000Δ*katA*Δ*ahpC*, which is unable to scavenge exogenously or endogenously produced H_2_O_2_, showed increased ROS production as compared to strain SH1000 (Fig. 8). Similarly, a higher fluorescence value per cell was obtained for the *dacA_G206S_* mutant as compared to WT and *gdpP* mutant cells (Fig. 8). These data suggest that ROS production is also increased in cells with low c-di-AMP levels, which might in turn contribute to the observed growth defect of the *dacA* mutant under aerobic conditions.

**Figure 8.**
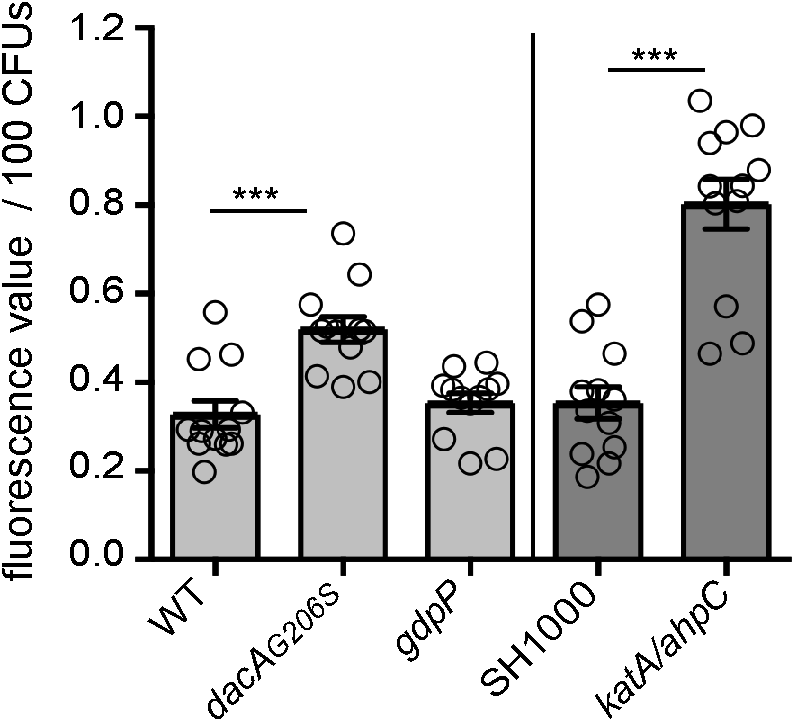
Reduced c-di-AMP levels lead to an increase in ROS production. Determination of endogenous ROS production in WT and mutant *S. aureus* strains. LAC* (WT), the isogenic *dacA_G206S_* and *gdpP* mutant strains as well as control strains SH1000 and SH1000Δ*katA*Δ*ahpC* were grown to mid-log phase in TSB medium and endogenous ROS production determined using the indicator dye 2’,7’-dichlorofluorescein diacetate used at a final concentration of 10 μM. Fluorescence values were measured at excitation and emission wavelengths of 485 nm and 538 nm, respectively. All fluorescent values were normalized for CFUs and the average fluorescence values and standard deviations per 100 CFUs from 12 biological replicates were plotted. Statistical analysis was performed in Prism (Graphpad) using for the comparison of the values obtained for the WT LAC* stain with those obtained for the isogenic *dacA_G206S_* and *gdpP* mutant strains (light grey bars) a Kruskal-Wallis test followed by a Dunn’s multiple comparison test. For the comparison of the values obtained for strains SH1000 and SH1000Δ*katA*Δ*ahpC* (medium grey bars), a Mann-Whitney test was used. Statistically significant differences are indicated by asterisks with p-values <0.001 indicated by a triple asterisk (***).

### DacA is dispensable for the growth of *S. aureus* under anaerobic conditions

Based on the reduced utilization of the main terminal oxidase Qox system in the *dacA* suppressor strains, we hypothesized that c-di-AMP production could become dispensable for growth under anaerobic conditions (Fig. 9A). To test this experimentally, the plating efficiencies of WT LAC* and the *dacA* mutant were determined by spreading culture aliquots on TSA plates and subsequently incubating them under aerobic and anaerobic conditions. This revealed that the *dacA* mutant grew similar to the WT strain under anaerobic conditions and had the same plating efficiency, indicating that c-di-AMP is dispensable when *S. aureus* is grown under anaerobic conditions (Fig. 9B and 9C). *S. aureus* can grow anaerobically by either glucose fermentation or, when nitrate (NO_3_^-^) is added to the medium, by NO_3_ respiration (Fig. 9A). The MQ pathway is required for nitrate respiration similar to aerobic respiration, except that a nitrate reductase and NO_3_^-^ are used as terminal acceptor in place of cytochromes and O2 as terminal acceptor (Fig. 9A and 9B). To determine whether c-di-AMP production is dispensable when *S. aureus* respires anaerobically on NO_3_, the WT and *dacA* mutant strains were plated on TSA plates supplemented with 20 mM KNO_3_ to allow NO_3_-dependent respiration to occur and the plates were subsequently incubated under either aerobic or anaerobic conditions. Under aerobic conditions, the *dacA* mutant showed the same plating defect on the nitrate-containing plates as observed on standard TSA plates (Fig. 9D). However, under anaerobic conditions the plating efficiencies of the WT and the *dacA* mutant were the same (Fig. 9D). Taken together, these experiments indicate that *dacA* is required during aerobic respiration through the Qox system but not for anaerobic nitrate respiration or anaerobic fermentation. Such findings have not been reported for other species, but it will be interesting to test in future studies how general these findings are with other *dacA* mutant bacteria.

**Figure 9.**
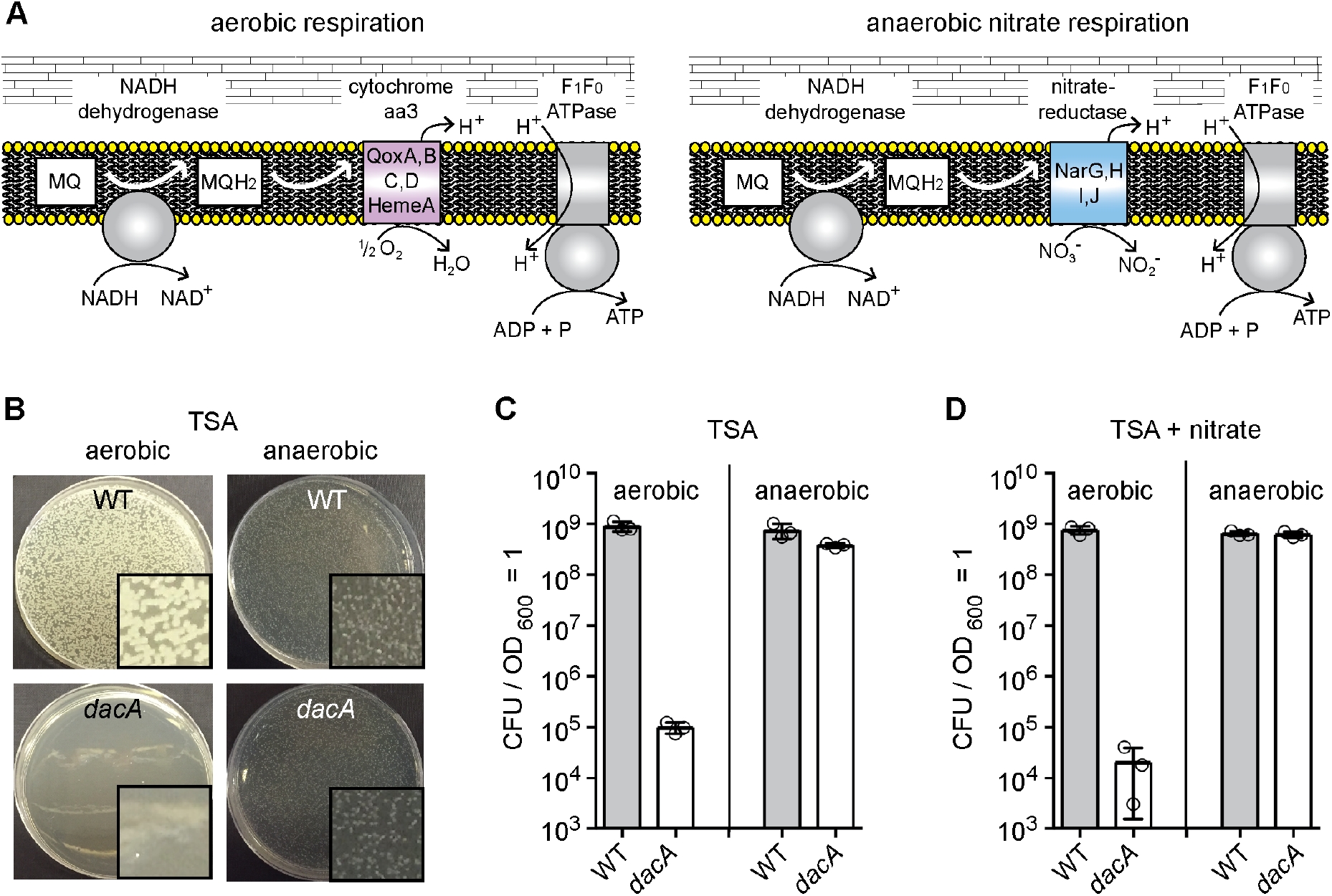
*dacA* is dispensable for the growth of *S. aureus* under anaerobic conditions. (A). Schematic overview of aerobic (left panel) versus anaerobic (right panel) respiration in *S. aureus*. Under aerobic conditions, oxygen is used as the terminal electron acceptor through the Qox system. In the absence of oxygen but in the presence of nitrate (NO_3_^-^), anaerobic respiration with nitrate as terminal electron acceptor through the nitrate reductase can occur. (B) Agar plate images. 10^-4^ dilutions of WT and *dacA* mutant cultures were plated on TSA plates, the plates incubated under aerobic or anaerobic conditions and images taken after overnight incubation at 37°C. Insets show representative magnified areas of the respective plate. (C-D) Plating efficiencies of WT and *dacA* mutant strains. Bacterial suspensions were plated on (C) TSA or (D) TSA plates containing 20 mM KNO_3_ and the plates incubated aerobically or anaerobically as stated. Average CFUs per ml culture per OD_600_ unit and SDs from three experiments were plotted.

### *S. aureus* strains with mutations that bypass the growth requirement of c-di-AMP remain hypersensitive to oxacillin

The level of c-di-AMP in different bacteria has been correlated with sensitivity to β-lactam antibiotics. Hence, a strain that completely lacks c-di-AMP and suppressors of the c-di-AMP phenotype allow this effect to be studied in more detail. In previous work, MRSA strains with increased levels of c-di-AMP (*gdpP* mutants) were found to have increased methicillin resistance, whereas strains with decreased levels of c-di-AMP displayed decreased methicillin resistance (15,20-22,51,52). Due to the previous unavailability of an *S. aureus* with a complete inactivation of *dacA*, no β-lactam resistance assays have been performed using an *S. aureus* strain completely lacking c-di-AMP. Standard minimal inhibitory concentration (MIC) assays with the β-lactam antibiotic oxacillin were performed in Müller Hinton medium (a rich medium) supplemented with 2% NaCl. While the colony size of the *S. aureus dacA* mutant was smaller as compared to a WT strain on this medium, the two strains had similar plating efficiencies on Müller Hinton 2% NaCl plates (Fig. 10A), likely owing to the addition of the extra 2% NaCl (0.2 M NaCl). This allowed us to perform oxacillin MIC assays with the *dacA* mutant strain completely lacking c-di-AMP. Similar to the low-level c-di-AMP *dacA_G206S_* control strain, the *dacA* strain was hyper-susceptible to oxacillin and showed a drastically reduced MIC towards this antibiotic (Fig. 9C). MIC assays were also performed with different *dacA* suppressor strains, which contained mutations in *ctaA, hemB, opuD, alsT* or *qoxB*. All strains remained hypersensitive to oxacillin, indicating that the growth improvement observed for these strains does not lead to a concomitant increase in oxacillin resistance (Fig. 10B). Taken together, these data show that a strain completely lacking c-di-AMP is hypersensitive to the β-lactam antibiotic oxacillin. Furthermore, commonly-acquired mutations that reverse the growth defect of a strain lacking c-di-AMP do not reverse the antibiotic sensitivity phenotype. This highlights that inhibiting DacA enzyme activity can potentially be used as strategy to re-sensitize MRSA strains to β-lactam antibiotics despite the ability of bacteria to acquire compensatory mutations that improves their growth in the absence of c-di-AMP production.

**Figure 10.**
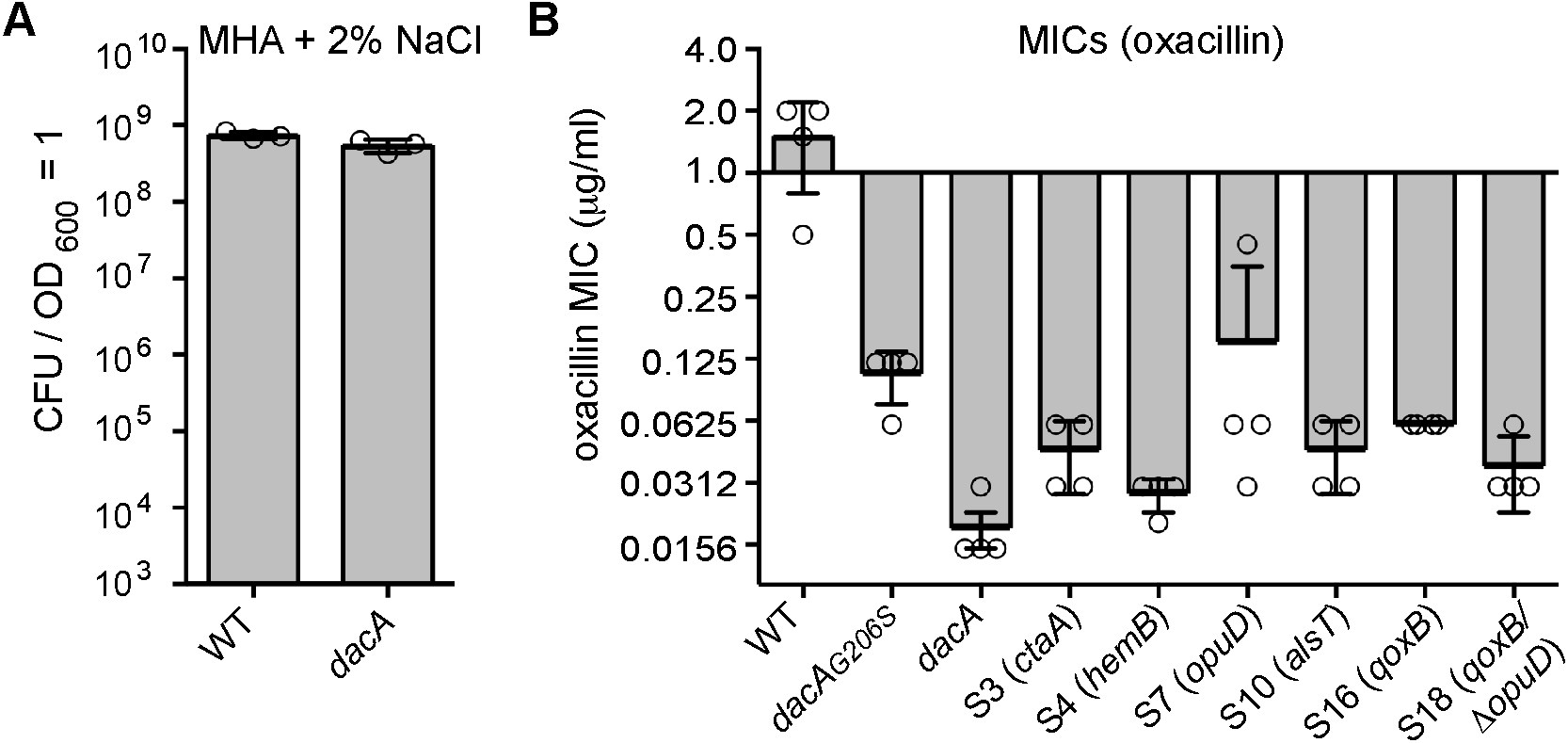
*dacA* mutant and suppressor strains have increased oxacillin sensitivity. (A) Plating efficiencies of WT and *dacA* mutant strains on Müller-Hinton agar (MHA) plates supplemented with 2% NaCl. Bacterial suspensions were prepared, plated on MHA plates supplemented with 2% NaCl and CFUs per ml culture per OD_600_ unit determined. The average values and SDs from three experiments were plotted. (B) Oxacillin MICs for WT and mutant *S. aureus* strains. Bacterial suspensions were prepared for WT LAC*, the low-level c-di-AMP *dacA_G206S_* strain, the *dacA* mutant and the indicated suppressor strains and spread on MHA plates with 2% NaCl. Next, oxacillin MIC Evaluator™ strips (ThermoScientific) were placed on the plates and the plates incubated for 24 hours at 35°C. The average MIC values and SDs from four independent experiments were plotted.

## DISCUSSION

In this study, we have investigated the requirement of c-di-AMP for the growth of *S. aureus*. c-di-AMP was previously shown to be required for the growth of *S. aureus* in rich medium (23), however, here we show that inactivation of the glycine betaine transporter OpuD, the predicted amino acid transporter AlsT or several proteins required for respiration can bypass the requirement of this signaling molecule for the growth of *S. aureus*. c-di-AMP is important for the growth of several other Gram-positive bacteria and bypass mutations that allow *L. monocytogenes* and *B. subtilis* to grow in the absence of c-di-AMP have been described (34,35,37). However, the types of mutations obtained in these two organisms were different. Whereas inactivation of the glycine betaine transporter Gbu and an oligopeptide uptake system Opp were described for *L. monocytogenes*, in *B. subtilis* inactivation of the high affinity potassium uptake system KimA (formally named YfhO) was observed. This indicates that in *L. monocytogenes* osmolyte or peptide uptake causes the detrimental effect on bacterial growth in the absence of c-di-AMP, whereas in *B. subtilis* it is excessive potassium uptake. However, it should be kept in mind that these suppressors screens were likely not performed to saturation. By expanding the screens, perhaps additional inactivating mutations in potassium transport systems, similar to what was observed in *B. subtilis*, could also be obtained in *L. monocytogenes* and *S. aureus*.

As part of this study, we show that inactivation of the *S. aureus* OpuD and AlsT transporters bypass the requirement of c-di-AMP for the growth in rich medium (Table 1 and Fig 4). AlsT is annotated as an amino acid transporter, and sometimes more specifically as an alanine symporter. However, no change in alanine uptake was detected when *alsT* was mutated, indicating that the encoded protein is likely not an alanine transporter, at least not under our growth conditions (Fig. 5E). Based on its homology and annotation, it is still likely that AlsT is responsible for the import of peptides or amino acids. This notion is supported by our observation that a *dacA* mutant suppressor strain with a mutation in *alsT* becomes insensitive to the addition of tryptone (as peptide source and hence amino acid source) to CDM medium, addition of which reduced the growth of the original *dacA* mutant (Fig. 5C). Furthermore, we show here that OpuD is likely the main glycine betaine uptake system in *S. aureus*, as its deletion leads to drastically reduced uptake of this osmolyte (Fig 5D). Taken together these data indicate that *S. aureus* can compensate for the lack of c-di-AMP using a similar mechanism to that described for *L. monocytogenes*, that is, by reducing the uptake of the osmolyte glycine betaine and likely also reducing the uptake of amino acids. It was also interesting to observe that the *opuD* and *alsT* suppressor strains were insensitive to the addition of both, glycine betaine and tryptone, but not either one or the other. This suggests a connection between the glycine betaine and amino acids uptake systems, perhaps through downstream metabolic activities in the cell that lead to the accumulation of the same toxic intermediate or depletion of an essential metabolite. While in *L. monocytogenes* it has been suggested that glycine betaine cannot be produced within the cell and must be taken up from an external source (53), a recent study revealed a flux of carbons from the amino acids glutamate, proline and arginine into betaine in *S. aureus* (36). This highlights a further metabolic link between amino acids and betaine production in *S. aureus*.

While inactivation of a glycine betaine transporter was observed in *S. aureus* and *L. monocytogenes* to bypass the need of c-di-AMP for their growth, it should be noted that the *S. aureus* OpuD and the *L. monocytogenes* Gbu glycine betaine transporter belong to different transporter classes. Whereas OpuD is a single protein transporter of the betaine/choline/carnitine (BCCT) transporter family, Gbu is a three-component ATP binding cassette (ABC) glycine betaine transporter (54), which is more closely related to the OpuC osmolyte uptake systems, previously described as a c-diAMP-regulated carnitine uptake system in *L. monocytogenes* and *S. aureus* (26,55). Interestingly, *L. monocytogenes* encodes a second glycine betaine transporter, BetL (Lmo02093 in strain EGDe) (56), which is more closely related to the OpuD transporter in *S. aureus*. However, no bypass mutations inactivating the BetL transporter have been found in *L. monocytogenes*. The reason for this might be that Gbu and not BetL is the main glycine betaine transporter in *L. monocytogenes*, at least under a number of different growth conditions (53). In our study, we also show that OpuD is likely the main glycine betaine transporter in *S. aureus* (Fig 4D).

We did not identify any mutations in the Opp peptide transport system, which was the second type of transporter identified in the *L. monocytogenes* suppressor screen. Homologous transport systems are present in *S. aureus*; however, in contrast to *L. monocytogenes*, which encodes only for a single Opp system, four different Opp transport systems are present in *S. aureus*, which mediate the uptake of peptides (57,58). While the four Opp transporters do not have completely overlapping functions, their activities might still be redundant enough so that inactivation of a single transporter is not sufficient to bypass the need of c-di-AMP for the growth of *S. aureus* on rich medium. Mutations in multiple systems might be required to compensate for the lack of c-di-AMP, which we were unlikely to obtained in our screen. Further pointing towards a role for c-di-AMP in the regulation of the osmolarity in the cell is our finding that bacteria producing low levels of c-di-AMP, such as observed in the *S. aureus dacA_G206S_* strain, are larger in size. Based on our cell diameter measurements, the cell volume of *dacA_G206S_* mutant bacteria is nearly twice that of a WT cell (Fig. 1C and 1D). In agreement with what has been proposed for *L. monocytogenes* (35), we envision that in a low c-di-AMP level strain, or in a strain completely lacking c-di-AMP, the increased uptake of osmolytes or accumulation of amino acids, leads to an increase in the intracellular osmotic pressure and thereby resulting in the observed increase in cell size. Consistent with this, we show here that increasing the external osmolarity through the addition of NaCl or KCl can rescue the growth defect of the c-di-AMP null strain in rich medium. This is in contrast to the observation that a reduction of potassium levels in the medium can rescue the growth defect in *B. subtilis* (37). Consistent with this, and perhaps indicating that excessive potassium transport is not the reason for the growth defect of *S. aureus* strains lacking the ability to produce c-di-AMP, we did not obtain suppressor mutations in genes coding for the constitutively expressed Ktr or conditionally-expressed Kdp potassium transport systems in *S. aureus*. However, the caveat that our suppressor screen was not performed to saturation needs to be kept in mind.

Perhaps one of the most interesting findings presented in this work, are our data showing that *S. aureus* can grow in the absence of c-di-AMP under anaerobic conditions (Fig 3, 6 and 9). The detrimental effect caused by the absence of c-di-AMP in rich medium observed during aerobic respiration through the main terminal oxidase system Qox is not detected during anaerobic growth, even if bacteria respire anaerobically when nitrate is provided as a terminal electron acceptor (Fig. 9). As part of our suppressor screen we identified mutations in multiple components of the respiratory chain, including proteins required for the synthesis of menaquinone, heme and cytochrome aa3, the main terminal oxidase used by *S. aureus* under high oxygen concentrations. During aerobic respiration, bacteria produce endogenous ROS, which at elevated levels can cause DNA damage as well as lipid and protein oxidation (48). We were not able to measure endogenous ROS production for the *dacA* mutant strain since this strain cannot be propagated under aerobic conditions in TSB medium. However, using a fluorescence dye indicator, we found that the low-level c-di-AMP *dacA_G206S_* mutant strain gave an increased fluorescent signal per cell indicative of increased endogenous ROS production at low cellular c-di-AMP levels (Fig. 8). We would expect that in the complete absence of c-di-AMP the endogenous ROS production might be even higher and the cellular damage caused through its production could contribute the observed growth inhibition of a *dacA* mutant strain under aerobic conditions.

Interestingly, our results indicate that aerobic respiration through the Cyd system and respiration under anaerobic conditions, using nitrate as terminal acceptor, is not detrimental in the absence of c-di-AMP. This indicates that it is not respiration *per se* but growth under aerobic respiration conditions and utilization of the proton-pumping Qox system is toxic to bacteria in the absence of c-di-AMP and hence excessive endogenous ROS production in the absence of c-di-AMP alone could not explain the observed growth defect in TSB medium. A large number of transcriptional and metabolic changes occur when bacteria are grown under anaerobic conditions (6). Notably, TCA cycle activity is reduced in bacteria grown under anaerobic conditions (6). Reduced TCA cycle activity is also seen under aerobic conditions in *S. aureus hemB* or *menD* mutants, the latter of which is unable to produce menaquinones (59-61) as well as a *B. subtilis qox* mutant strains (62). This is of particular interest in connection with the c-di-AMP signaling pathway, as in *L. monocytogenes* accumulation of high levels of citrate, a key TCA cycle intermediate, has been shown to contribute to the growth inhibition caused by the absence of c-di-AMP (35). An *L. monocytogenes dacA* mutant also lacking *citZ*, which encodes the citrate synthase, a key TCA cycle enzyme, was viable in rich medium (35). The TCA cycle is also a key cellular process that is required for the production and interconversion of many different amino acids (36). One amino acid that has a particularly critical role in TCA cycle function is glutamate, which also plays a key role in the osmotic regulation in bacterial cells. Hence, the suppressor mutations we obtained in the AlsT and OpuD transporters might not only allow bacterial cells to regain the ability to adjust their osmotic balance, but also indirectly impact the TCA cycle activity, through the crosstalk between osmolytes, amino acids and TCA cycle activity.

Finally, we show that the *dacA* mutant MRSA strain used in this study, which is unable to produce c-di-AMP, is hyper-sensitive to the β-lactam antibiotic oxacillin (Fig. 10). Mechanistically, how c-di-AMP impacts β-lactam resistance is currently not known. Recently it has been proposed that c-di-AMP levels might not lead to changes in the peptidoglycan structure *per se*, but that c-di-AMP impacts the sensitivity of bacteria to β-lactam antibiotics by modulating the turgor pressure of the cell, a physical, not a structural variable (63). In future studies, it will be interesting to investigate if resistance to cell wall-targeting antibiotics other than β-lactam antibiotics is impacted by c-di-AMP levels, as one might expect if the sensitivity is linked to the intracellular turgor pressure. However, it is of note that the *dacA* suppressor strain inactivated for OpuD remained hypersensitive to β-lactam antibiotics. This suppressor strain showed reduced glycine-betaine uptake activity and hence we would expect might also have a reduced turgor pressure. Hence the reason for the β-lactam hypersensitivity of *dacA* mutant strains might be multifactorial and not solely due to an increase in cellular turgor pressure.

The pathways regulated by the c-di-AMP system and the impact of c-di-AMP on bacterial growth and physiology are clearly very complex. As part of this study we have provided novel information on the requirement of c-di-AMP for the growth of *S. aureus*, but additional work is needed in order to uncover the mechanistic details behind this. Our work opens up a number of interesting avenues for further research; in particular, it will be of interest to test if other bacteria can grow under anaerobic conditions in the absence of c-di-AMP, similar to *S. aureus*. Our work also highlights how difficult the definition of essential genes is, in particular for nutritionally related genes. We show here that not only growth conditions but also other physical parameters such as the availability of oxygen can heavily influence the essentiality of genes.

## EXPERIMENTAL PROCEDURES

### Bacterial strains and culture conditions

Bacterial strains used in this study are listed in Table 2. *E. coli* strains were grown in Lysogeny Broth (LB) broth or agar and *S. aureus* strains in Tryptic Soy Broth (TSB), Tryptic Soy Agar (TSA), or Chemically Defined Medium (CDM) adjusted to pH 7.2. The CDM was prepared using components based on two previously described recipes (64,65) and contained: KCl 3 g/L, NaCl 9.5 g/L, MgSO_4_ 7H_2_O 1.3 g/L, (NH_4_)_2_SO_4_ 4 g/L, Tris 12.1 g/L, Glucose 5 g/L, L-Arg 50 mg/L, L-Pro 10 mg/L, L-Glu 100 mg/L, L-Val 80 mg/L, L-Thr 30 mg/L, L-Phe 40 mg/L, L-Leu 90 mg/L, L-Gly 50 mg/L, L-Ser 30 mg/L, L-Asp 90 mg/L, L-Lys 50 mg/L, L-Ala 60 mg/L, L-Trp 10 mg/L, L-Met 10 mg/L, L-His 20 mg/L, L-Ile 30 mg/L, L-Tyr 50 mg/L, L-Cysteine 20 mg/L, Biotin 0.1 mg/L, Thiamine 2 mg/L, Nicotinic acid 2 mg/L, Calcium pantothenate 2 mg/L, CaCl_2_ 2H_2_O 22 mg/L, KH_2_PO_4_ 140 mg/L, FeSO_4_ 7H_2_O 6 mg/L, MnSO_4_ 4H_2_O 10 mg/L, Citric acid 6 mg/L). For the alanine uptake assays, bacteria were grown in CDM with ½ the L-Ala concentration (30 mg/L). Where indicated, the TSA plates were supplemented with 400, 800, 1200, 1600 or 2000 mM NaCl or KCl, 20 mM KNO_3_ or 10 μM hemin. Cation-adjusted Müller-Hinton agar plates supplemented with 2% NaCl were used to determine the oxacillin minimal inhibitory concentration (MIC) for the different *S. aureus* strains. When needed, antibiotics and/or inducers were added to the media at the following concentration: 200 ng/ml anhydrotetracyline (Atet), 90 or 200 μg/ml Kanamycin (Kan), 2 μg/ml Tetracycline (Tet), 10 μg/ml Erythromycin (Erm), 7.5 μg/ml Chloramphenicol (Cam), 100 μg/ml Ampicillin (Amp). Bacteria were grown for 16 to 24 h aerobically and where specified anaerobically, by incubating plates in an anaerobic cabinet (Don Whitley Scientific) in an atmosphere of 10% CO_2_, 10% H_2_ and 80% N_2_.

**Table 2:**
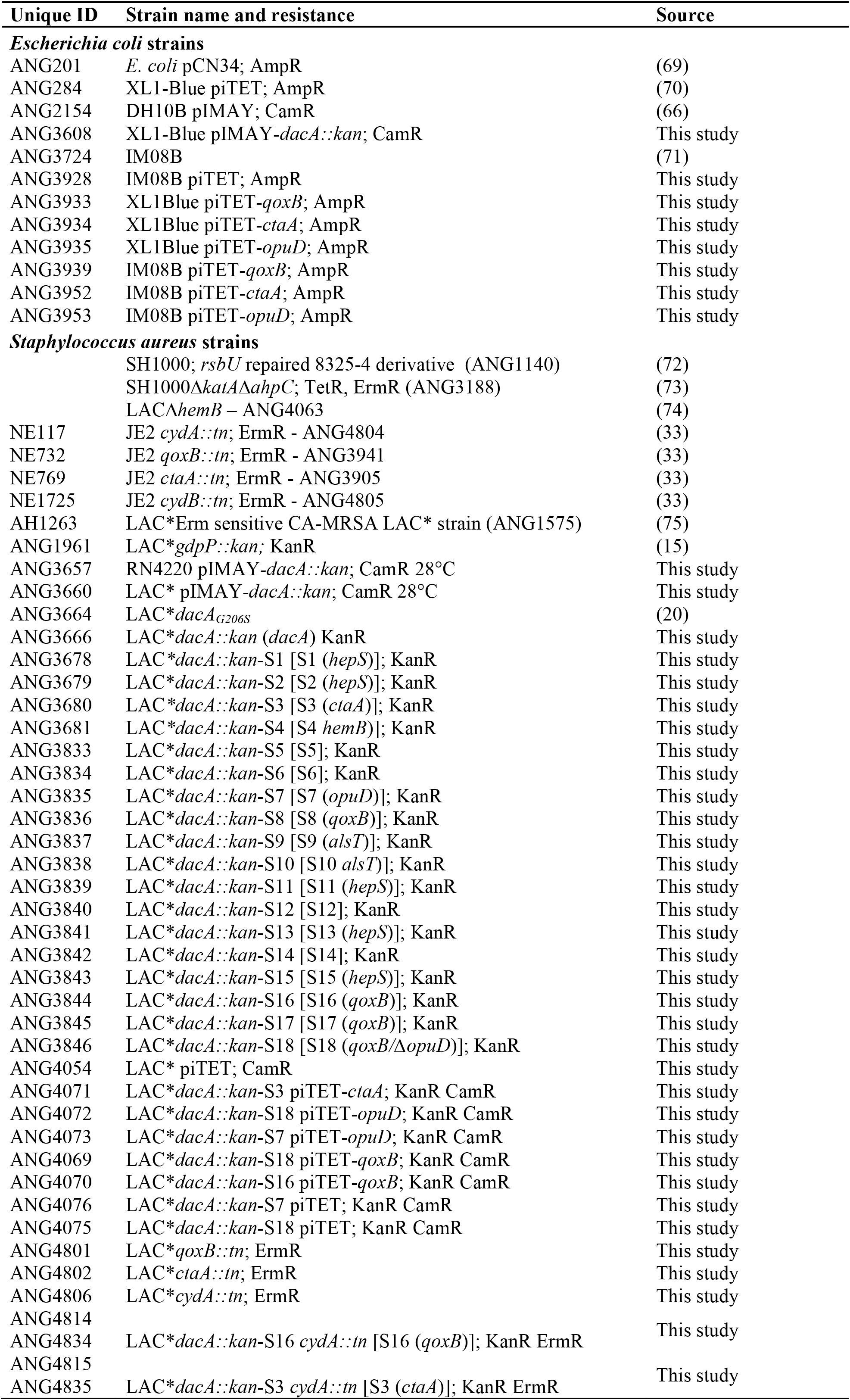
Bacterial strains used in this study

### Bacterial strain construction

Primers used for plasmid and strain construction are listed in Table 3. For construction of the *dacA* mutant *S. aureus* strain LAC\**dacA::kan* (ANG3666) in which the *dacA* gene is replaced with the *aph3* gene conferring kanamycin resistance, the allelic exchange vector *pIMAY-dacA::kan* was produced. To this end, approximately 1 kb up and downstream regions of *dacA* were amplified using LAC* chromosomal DNA and primer pairs ANG2105/ANG1191 and ANG1194/ANG2106, respectively. The *aph3* gene was amplified from plasmid pCN34 using primers ANG2105/ANG1191. The *aph3* gene and the downstream *dacA* fragment were fused by splicing overlap extension (SOE) PCR using primers ANG1192/ANG2106 and the upstream *dacA* fragment subsequently fused by SOE PCR using primers ANG2105/ANG2106. The product was cut with EcoRI and XmaI and cloned into vector pIMAY that had been cut with the same enzymes. *pIMAY-dacA::kan* was initially recovered in *E. coli* strain XL1-Blue yielding strain ANG3608. The plasmid was subsequently moved through *S. aureus* strain RN2440 (ANG3608) into *S. aureus* strain LAC* and colonies recovered at 28°C yielding strain ANG3660. Allelic exchange was performed using a standard procedure (66) with the exception that the plasmid resolution step at 28°C was performed in CDM and bacteria were plated in the final step on CDM plates supplemented initially with 200 μg/ml kanamycin. This resulted in the construction of strain LAC\**dacA::kan* (or short *dacA)*. The replacement of the *dacA* gene with the *aph3* gene was confirmed by sequencing and the strain was propagated in CDM supplemented with 90 μg/ml kanamycin.

**Table 3:**
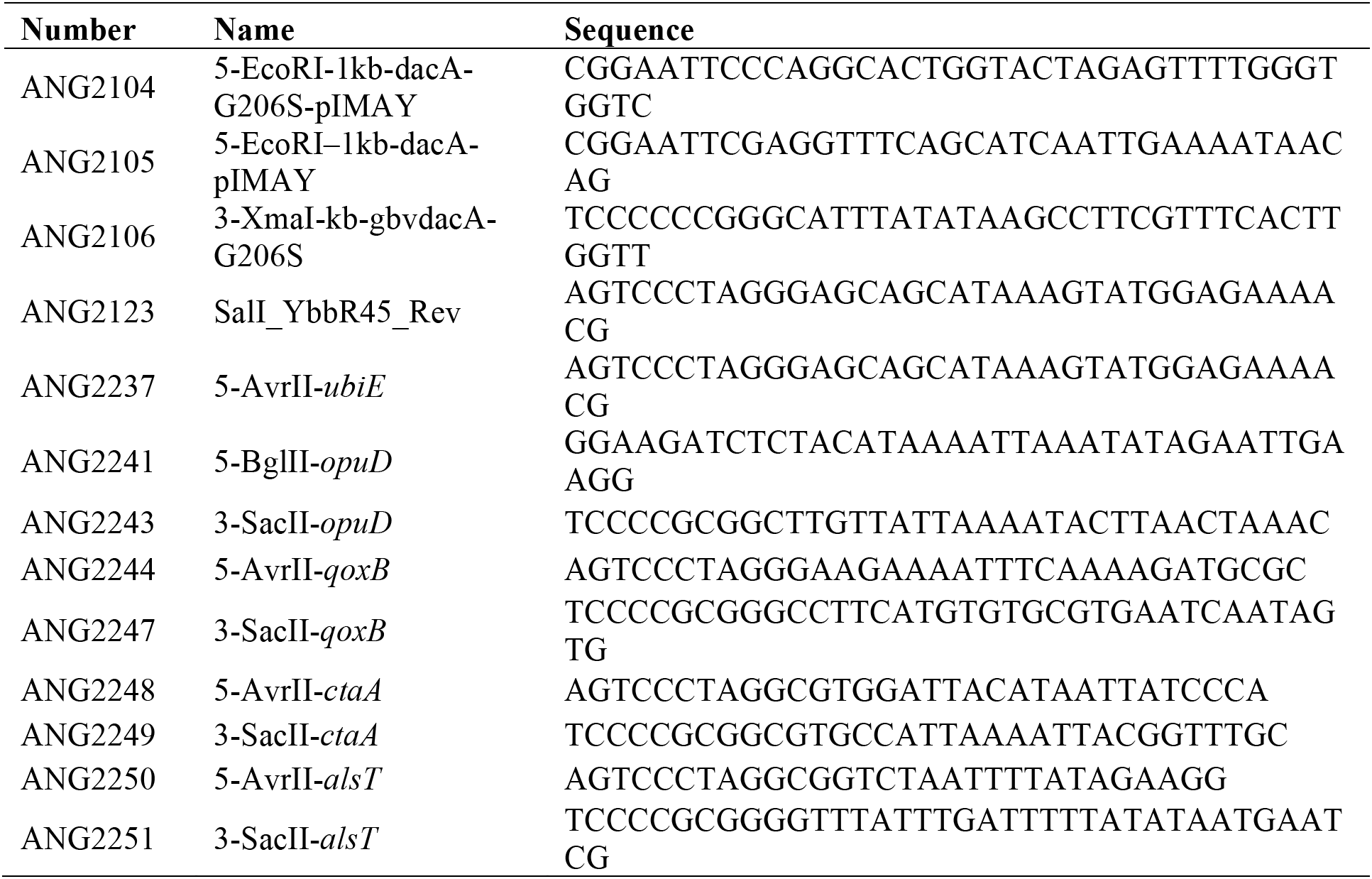
Primers used in this study.

For complementation analysis, single copy integration plasmids piTET-qoxB, piTET-ctaA and piTET-opuD were constructed, which allow for Atet inducible gene expression in *S. aureus*. To this end, the genes of interest were amplified from LAC* chromosomal DNA using primer pairs ANG2244/ANG2247 (*qoxB*), ANG2248/ANG2249 (*ctaA*) and ANG2241/ANG2243 (*opuD*) and the resulting products were then digested with AvrII (BglII for the *opuD*) and SacII and ligated with piTET that had been digested with the same enzymes. Plasmids piTET-ctaA, piTET-opuD and piTET-qoxB were individually recovered in *E. coli* strain XL1-Blue, yielding strains ANG3933-ANG3935. Next the plasmids were shuttled through *E. coli* strain IM08B (yielding strains ANG3939, ANG3952 and ANG3953) and finally introduced by electroporation into the appropriate LAC\**dacA::kan* suppressor strains. Transformants were recovered on CDM plates under aerobic conditions or on TSA plates under anaerobic conditions, which yielded the complementation strains LAC* *dacA:: kan-S3* piTET-ctaA (ANG4071), LAC* *dacA::kan-S7* piTET-opuD (ANG4073), LAC\**dacA::kan-S16* piTET-qoxB (ANG4070), and LAC\**dacA::kan-S18* piTET-opuD (ANG4072). Correct plasmid integration into the *geh* locus was confirmed by PCR and the sequences of all plasmid inserts were confirmed by fluorescent automated sequencing. As controls, the empty piTET vector was also introduced by electroporation in two of the *dacA* suppressor strains, yielding strains LAC\**dacA::kan-S7* piTET (ANG4076) and LAC\**dacA::kan-S18* piTET (ANG4075). *S. aureus* strains from the Nebraska transposon library (33) with transposon insertions in *qoxB* (NE732), *ctaA* (NE769) and *cydA* (NE117) were used as part of this study. The transposon insertions in the respective genes were confirmed by PCR and sequencing. Subsequently the transposons were moved into the LAC* strain background by transduction using phage 85 yielding strains LAC* *qoxB::tn* (ANG4081), LAC\**ctaA::tn* (ANG4082) and LAC\**cydA::tn* (ANG4806). The *cydA* transposon mutation was also moved into the *dacA* suppressor strains S3 and S4 yielding strains LAC\**dacA::kan-S3 cydA::tn* (ANG4815 and ANG4835) and LAC* *dacA::kan*-S16 *cydA::tn* (ANG4814 and ANG4834), respectively and transposon insertions in the appropriate gene confirmed by PCR.

### Bacterial growth curves

For the bacterial growth curve measurements (Figs. 4 and 6), the indicated *S. aureus* strains were grown overnight in TSB or CDM (strain LAC\**dacA::kan*) supplemented with the appropriate antibiotics. The overnight cultures were diluted in 50 ml fresh TSB as well as CDM (without antibiotics) to a starting OD_600_ of 0.01 and the cultures were incubated at 37°C with aeration. For the growth curve analyses using CDM supplemented with 1 mM glycine betaine (GB) or 1% tryptone, the different *S. aureus* strains were streaked onto CDM agar plates with the appropriate antibiotics, and the plates incubated overnight at 37°C. The next day, patches of bacteria were removed from the plate and suspended in 1 ml PBS pH 7.4 buffer. The culture suspensions were diluted in 50 ml of CDM, CDM 1 mM glycine betaine, or CDM 1% tryptone to a starting OD_600_ of 0.01 and incubated at 37°C with aeration. For all growth curves, OD600 values were measured every two hours and three independent experiments were performed.

### Determination of plating efficiencies

*S. aureus* strains were streaked onto CDM agar plates containing the appropriate antibiotics, and the plates incubated overnight at 37°C. The next day, patches of bacteria were removed from the plate and suspended in 1 ml PBS pH 7.4 buffer. The suspensions were normalized to an OD_600_ of 2. Serial dilutions from 10^-2^ to 10^-6^ were made and 100 μl plated onto TSA, TSA 200 ng/ml Atet, TSA 20 mM potassium nitrate (KNO_3_), TSA 10 μM hemin, or CDM plates, as indicated. The plates were incubated under aerobic or anaerobic conditions for 16 to 20 h. The colony forming units (CFUs) per ml suspension per OD_600_ unit were determined and plotted. The experiments were performed with at least three biological and two technical replicates. To determine the plating efficiencies of the WT LAC* and *dacA* mutant strain on TSA plates supplemented with increasing NaCl and KCl concentrations, bacteria were scraped from CDM plates and suspended in 1 ml PBS pH 7.4 buffer and the suspensions normalized to an OD_600_ of 2. Tenfold serial dilutions were prepared, and 5 μl of the 10°-10^-7^ dilutions spotted onto TSA or TSA plates supplemented with NaCl or KCl at the indicated concentrations. The plates were incubated for 20 h at 37°C and colonies enumerated. The colony forming units (CFUs) per ml suspension per OD_600_ unit were calculated and the average and standard deviation of three independent experiments plotted.

### Whole genome sequencing

The genome sequencing for suppressor strains ANG3678-3681 (S1-S4) was performed by MicrobesNG (http://microbesng.uk) using an Illumina HiSeq platform and a 250 bp paired end read kit. Suppressor strains ANG3833-3846 (S6-S18) were sequenced at the Department of Microbiology and Immunobiology at Harvard Medical School using an Illumina MiSeq platform and a 150 bp paired end read kit. The CLC Genomics Workbench Software was used for genome sequence analysis. As reference genome, a contig produced for the WT LAC* strain was used. This contig was produced as part of a previous study (20) by mapping Illumina reads onto the closely related strain USA300 FPR3757 (RefSeq accession number NC_007793.1) genome sequence and transferring its annotation. The Illumina short read sequences from the different suppressor strains were mapped onto the assembled LAC* sequence, and good quality and high frequency (>65%) base changes and small deletions and insertions were identified using the CLC Genomics Workbench Software. Large deletions were identified in a second step by specifically searching for zero coverage regions. The genome sequence data listing the base changes in the suppressor strains as compared to LAC* are presented in Table 1. Sequence variations detected by the whole genome sequence analysis in *hepS*, *qoxB, ctaA, opuD, hemB, alsT* were subsequently confirmed by fluorescence automated sequencing of PCR products of the respective genome regions. The Illumina reads for the LAC* strain have been deposited in the European Nucleotide Archive under the study accession numbers PRJEB14759 as part of a previous study (20). The Illumina short reads for the *dacA* suppressor strains (S1-S4 and S6-S18) were deposited under accession number PRJEB22312.

### LC-MS/MS detection of c-di-AMP in *S. aureus* extracts

An LC-MS/MS analysis was used for the detection of c-di-AMP in *S. aureus* extracts. Three independent extracts were prepared for the LAC*\*dacA::kan*-S1, LAC\**dacA::kan-S3* and LAC\**dacA::kan-S4* suppressor strain. The sample preparation and LC-MS/MS analysis was performed in the same manner and at the same time as the WT LAC* and LAC\**dacA_G206S_* described in Bowman *et al*. (20). The data from these two control strains are shown as controls in the graph. c-di-AMP could not be detected in any of the extracts derived from the suppressor strains.

### Membrane potential measurements

The membrane potential of different *S. aureus* strains was measured using the 3, 3’-diethyloxacarbocyanine iodide [DiOC_2_(3)] dye and a previously described fluorescence activated cell sorter (FACS)-based method (42). At a high dye concentration, the emitted green fluorescence is dependent on cell size, while the emitted red fluorescence is dependent on both cell size and membrane potential. Therefore, the ratio of red/green fluorescence gives a largely cell-size independent measure of the membrane potential (42). For the measurements, *S. aureus* strains LAC*, LAC\**gdpP::kan* and LAC* dacA_G206S_ were grown overnight at 37°C with aeration in 5 ml TSB medium. The next day, bacteria from 1 ml sample with an OD_600_ of 1 were harvested by centrifugation at 17,000 *xg* and the cells washed 3 times with 1 ml of staining buffer (0.06 M Na_2_HPO_4_, 0.06 M NaH_2_PO4, 5mM KCl, 130 mM NaCl, 1.3 mM CaCl_2_, 0.5 mM MgCl_2_, 10 mM glucose). After the final wash step, the cells were diluted to an OD_600_ of 0.2 in staining buffer. 100 μl of these cell suspensions were added to FACS tubes containing 890 μl staining buffer and 10 μl of 3 mM DiOC_2_(3) dye (giving final OD_600_ = 0.02 and 30 μM dye concentration). The mixtures were incubated at room temperature for 10 minutes. As control, 7.5 μl of 2 mM carbonyl cyanide m-chlorophenylhydrazone (CCCP) (15 μM final concentration) was added to a second set of samples containing the cells and the DiOC_2_(3) dye. CCCP collapses the membrane potential and bacteria will show green fluorescence, but will appear with only a background level of red fluorescence, which aids in the identification and gating on the bacterial cell population during the FACS analysis. All samples were acquired using a Becton Dickinson two-laser, four color FACSCalibur cytometer and the Cell Quest Pro software analysis software. Fluorophores were excited with a 488 nm or 633nm laser. Green and red fluorescence intensities were detected in FL-1 (530/30 nm) and FL-2 (585/42 nm) channels, respectively. 10,000 gated cells were recorded for each sample with the mean fluorescence intensity of each event being measured at the height of its emission peak. The ratio of red / 10 *x* green fluorescence was determined using the FlowJo Version 7 (Tree Star) software and histograms of cell counts versus fluorescence ratio were produced from these data. From the histograms, the mean red / 10 *x* green fluorescence values were also determined using the FlowJo software and averages and SDs from six biological replicates were plotted.

### Microscopic analysis and bacterial cell size determination

*S. aureus* strains LAC*, LAC* *gdpP::kan* and LAC* dacA_G206S_ were grown overnight at 37°C in TSB. Bacterial cultures were normalized to an OD_600_ of 1 and 100 μl of these cultures stained for 20 min at 37°C with Vancomycin-BODIPY FL at a final concentration of 2 μg/ml. The stained bacteria were collected by centrifugation for 1 min at 17,000 *xg*. The cells were washed with 100 μl PBS pH 7.4, centrifuged again and finally suspended in 100 μl PBS pH 7.4. 1.5 μl sample was spotted onto a thin 1.5% agarose gel patch prepared in ddH_2_O and imaged at 1000 × magnification using an Axio Imager.A2 Zeiss microscope equipped with a GFP filter set. Images were acquired using the ZEN 2012 (blue edition) software. Staining *S. aureus* cells with Vancomycin-BODIPY FL allows one to determine the outline of bacterial cells as well as to distinguish between dividing and non-dividing bacterial cells, which do not have dots or a line at mid cell. The bacterial cell diameters were determined by measuring the pixel width of the bacterial cells using the Image J software. For these measurements, only non-dividing cells, cells without any obvious fluorescent dots or lines at the mid-plane, were used. The length in pixels was subsequently converted to μm using a conversion scale provided in the ZEN 2012 (blue edition) software. Images of cells from three biological replicates were taken and 50 cells were measured per biological replicate (150 cells in total per strain). The mean and standard deviation for 150 cells per strain were plotted.

### Glycine betaine and alanine uptake assay

Uptake assays were essentially conducted as described in Schuster *et al*, (26). The different *S. aureus* strains were streaked on CDM plates with appropriate antibiotics and the plates incubated aerobically overnight at 37°C. Bacteria were subsequently scraped off from the plates and suspended in 1 ml PBS pH 7.4 buffer and the OD_600_ determined. For glycine-betaine uptake assays, 50 ml of standard CDM (without antibiotics but containing 200 ng/ml Atet) and for the alanine uptake assays CDM medium with ½ the amount of alanine was inoculated with the appropriate bacterial suspensions to give a starting OD600 of 0.01. The cultures were grown with shaking at 37°C to an OD_600_ between 0.5 and 0.9 and an OD_600_ equivalent of 8 was harvested by centrifugation for 10 min at 19,000 *xg* and 4°C. Supernatants were carefully discarded and the pellet suspended by swirling in 2 ml of CDM (glycine betaine uptake) or CDM ½ alanine (alanine uptake). The OD_600_ of the cell suspensions were measured and the cells diluted to an OD_600_ of approximately 1. The OD 600 was re-measured after the dilution and this reading used for normalization purposes. 450 μl of these cell suspensions were aliquoted into 50 ml conical tubes and 100 μl used to measure the background radiation, by filtering the cell onto nitrocellulose filter and then washing them with 16 ml of CDM or PBS (alanine uptake). Then, 4.8 μl of glycine-1-^14^C (Hartmann Analytic) or D/L-alanine-(1)-^14^C (Hartmann Analytic) was added to the remaining 350 μl sample yielding a final concentration of 25 μM radiolabeled compound. 100 μl aliquots were filtered at 0, 3 and 6 minutes and the filters washed with 2 × 16 ml of CDM or PBS pH 7.4 (alanine uptake). The filters were subsequently dissolved in 9 ml of scintillation cocktail Filter Count (Perkin Elmer) and the radioactivity measured in counts per minute (CPM) using a Wallac 1409 DSA liquid scintillation counter. CPMs of each sample were then normalized by the OD_600_ of the final cell suspension and the means and standard deviations of the CPM / ml OD_600_ = 1 of for four independent experiments plotted.

### Determination of oxygen consumption rates

Oxygen consumption rates were determined using a Clark-type dissolved oxygen electrode (Rank Brothers Ltd) and a previously described method with some modifications (67). Briefly, the indicated *S. aureus* strains were grown overnight at 37°C in TSB. The next day, the cultures were diluted to an OD_600_ of 0.01 (or 0.05 for strain LACΔhemB, LAC\**dacA::kan-S3 cydA::tn* (ANG4815 and ANG4835) and LAC\**dacA::kan-S16 cydA::tn* (ANG4814 and ANG4834) and grown for 3 hours to mid-log phase (OD600 of 0.2 to 0.9). Bacteria from 30-50 ml culture aliquots were harvested by centrifugation for 10 min at 9,000 *x g* and washed twice with 20 ml PBS pH 7.4 buffer and then suspended in 100-200 μl of PBS pH 7.4 buffer. Next, the cultures were set to an OD_600_ of 50 in PBS pH 7.4 buffer and kept on ice for a few minutes until the oxygen consumption experiment was started. Prior to measuring the bacterial respiration rates, the maximal response rate of the oxygen electrode was determined each day as follows: the electrode was equilibrated with 3 ml PBS pH 7.4 buffer and the chamber maintained at 37°C using a circulating water bath. The signal (maximal reading) for the 3 ml buffer assumed to contain 720 nmol oxygen was recoded for 1 to 2 min using a chart recorder or data logger. Next, all oxygen in the buffer was consumed by the addition of a few mg sodium hydrosulphite and the signal followed for a few minutes until it plateaued, yielding the minimal reading and hence the maximal response range following the consumption of the 720 nmol oxygen in the buffer. For measuring the oxygen consumption rates of the different *S. aureus* strains, the oxygen electrode was again equilibrated with 3 ml PBS pH 7.4 buffer and the chamber maintained at 37°C. Next, 50 μl of the washed bacterial suspensions with an OD_600_ of 50 were injected into the chamber, and the response signal recoded for 2 min to determine a base-line oxygen consumption. Then, 30 μl of 0.1 M glucose solution was added as previously described for the determination of oxygen consumption rates in *S. aureus* (68), and the signal subsequently recorded for a further 3 min to 5 min. The oxygen consumption rate associated with the addition of glucose was determined from the slope of the response curve. The experiment was performed with 3 biological replicates, and the average and standard deviation of the oxygen consumption rate per minute per ml culture of OD_600_ = 1 following the addition of glucose plotted.

### Determination of endogenous ROS production

Endogenous ROS production was determined using the 2’,7’-dichlorofluorescein diacetate indicator dye and a previously described method with some modifications (49,50). *S. aureus* strains LAC*, LAC* *gdpP::kan* and LAC* *dacA_G206S_* as well as the *S. aureus* control strains SH1000 and SH1000Δ*katA*Δ*ahpC*, the latter of which is unable to scavenge exogenously or endogenously produced H_2_O_2_, were grown overnight at 37°C in TSB. The next day, the cultures were diluted to an OD_600_ of 0.01 or 0.025 for strain SH1000Δ*katA*Δ*ahpC* and grown for 3 hours to midlog phase (OD_600_ of 0.3 to 1). Bacteria from 2-4 ml culture aliquots were harvested by centrifugation and washed twice with 1 ml PBS pH 7.4 buffer. Next, the cultures were set to an OD_600_ of 0.5 and 100 μl dispensed into a 96-well plate and mixed with 100 μl of 20 μM 2’,7’-dichlorofluorescein diacetate in PBS buffer to give a final dye concentration of 10 μM. The plates were incubated with shaking for 2h at 37°C and fluorescence values subsequently measured in a fluorescent plate reader using excitation and emission wavelengths of 485 nm and 538 nm, respectively. Following the measurements, the number of CFU per well was determined by plating appropriate dilutions on TSA plates. The background fluorescence value obtained for well containing a 10 μM dye solution in PBS was subtracted from the fluorescence values obtained for wells containing the bacterial suspensions. All fluorescent values were subsequently normalized for CFU counts. The experiment was performed with performed with 12 biological replicates, and the average fluorescence values per 100 CFUs were plotted along with the standard deviations.

### Determination of oxacillin minimal inhibitory concentrations (MICs)

WT LAC*, LAC* *dacA_G206S_*, and the LAC* *dacA::kan* suppressor strains were streaked onto CDM plates and the plates incubated overnight at 37°C. The next day, the bacteria were scraped off the plates, suspended in 1 ml of PBS and diluted to an OD600 of 0.1 and 250 μl spread onto cation adjusted Müller-Hinton agar plates containing 2% NaCl. Oxacillin M.I.C. Evaluator™ (ThermoScientific) strips were then placed onto the agar, and the plates were incubated at 35°C for 24 h and the MIC read on the strips. Four biological replicates were performed and the average values and standard deviations plotted.

## ACKNOWLEDGMENTS

This research was supported by the European Research Council grant 260371 and the Wellcome Trust grant 100289 to A.G.; M.S.Z. is supported by a Medical Research Council Centre for Molecular Bacteriology and Infection (MRC CMBI) studentship and C.F.S. by the German research foundation (DFG) grant SCHU 3159/1-1. The Illumina sequencing of the Suppressor strains S1-S4 was performed by MicrobesNG (http://www.microbesng.uk), which is supported by the BBSRC (grant number BB/L024209/1). We would like to thank Thomas Bernhardt and Jackson Buss for help with the Illumina sequencing of some of the suppressor strains and Eleni Karinou for construction of strains ANG3928 and ANG4054. The FACS experiment was performed at the Flow Cytometry Facility at Imperial College London and we would like to thank Jane Srivastava and Sophie Helaine for help with this analysis. We would like to thank Andrew Edwards for advice on ROS assays and we also acknowledge the assistance of the Research Core Unit Metabolomics at the Hannover Medical School for the LC-MS/MS experiment and c-di-AMP detection.

## AUTHOR CONTRUIBUTION

M.S.Z., C.F.S., L.B. H.D.W. and A.G. designed the experiments; M.S.Z., C.F.S. and Q.Z. performed the experiments; Q.Z. analyzed the microscopy data, C.F.S. analyzed the glycine betaine uptake assay data, L.B. analyzed the LC-MS/MS data; H.D.W. analyzed the oxygen consumption rate data, M.S.Z. and A.G. analyzed all data presented in the manuscript and wrote the manuscript. C.F.S. contributed to the writing of the manuscript and L.B. and Q.Z. edited the manuscript and all authors approved of the final connect in the manuscript.

## References

1. Kluytmans, J., van Belkum, A., and Verbrugh, H. (1997) Nasal carriage of *Staphylococcus aureus:* epidemiology, underlying mechanisms, and associated risks. Clin Microbiol Rev 10, 505–520

2. Fridkin, S. K., Hageman, J. C., Morrison, M., Sanza, L. T., Como-Sabetti, K., Jernigan, J. A., Harriman, K., Harrison, L. H., Lynfield, R., Farley, M. M., and Active Bacterial Core Surveillance Program of the Emerging Infections Program, N. (2005) Methicillin-resistant *Staphylococcus aureus* disease in three communities. N Engl J Med 352, 1436–1444

3. Francis, J. S., Doherty, M. C., Lopatin, U., Johnston, C. P., Sinha, G., Ross, T., Cai, M., Hansel, N. N., Perl, T., Ticehurst, J. R., Carroll, K., Thomas, D. L., Nuermberger, E., and Bartlett, J. G. (2005) Severe community-onset pneumonia in healthy adults caused by methicillin-resistant *Staphylococcus aureus* carrying the Panton-Valentine leukocidin genes. Clin Infect Dis 40, 100–107

4. Ferreira, M. T., Manso, A. S., Gaspar, P., Pinho, M. G., and Neves, A. R. (2013) Effect of oxygen on glucose metabolism: utilization of lactate in *Staphylococcus aureus* as revealed by in vivo NMR studies. PLoS One 8, e58277

5. Pagels, M., Fuchs, S., Pane-Farre, J., Kohler, C., Menschner, L., Hecker, M., McNamarra, P. J., Bauer, M. C., von Wachenfeldt, C., Liebeke, M., Lalk, M., Sander, G., von Eiff, C., Proctor, R. A., and Engelmann, S. (2010) Redox sensing by a Rex-family repressor is involved in the regulation of anaerobic gene expression in *Staphylococcus aureus*. MolMicrobiol 76, 1142–1161

6. Fuchs, S., Pane-Farre, J., Kohler, C., Hecker, M., and Engelmann, S. (2007) Anaerobic gene expression in *Staphylococcus aureus*. JBacteriol 189, 4275–4289

7. Tynecka, Z., Szczesniak, Z., Malm, A., and Los, R. (1999) Energy conservation in aerobically grown *Staphylococcus aureus*. Res Microbiol 150, 555–566

8. Collins, M. D., and Jones, D. (1981) Distribution of isoprenoid quinone structural types in bacteria and their taxonomic implication. Microbiol Rev 45, 316–354

9. Götz, F., and Mayer, S. (2013) Both Terminal Oxidases Contribute to Fitness and Virulence during Organ-Specific *Staphylococcus aureus* Colonization. Mbio 4

10. Hammer, N. D., Reniere, M. L., Cassat, J. E., Zhang, Y., Hirsch, A. O., Indriati Hood, M., and Skaar, E. P. (2013) Two heme-dependent terminal oxidases power *Staphylococcus aureus* organ-specific colonization of the vertebrate host. MBio 4

11. Römling, U. (2008) Great times for small molecules: c-di-AMP, a second messenger candidate in Bacteria and Archaea. Sci Signal 1, pe39

12. Hengge, R. (2009) Principles of c-di-GMP signalling in bacteria. Nat Rev Microbiol 7, 263–273

13. Witte, G., Hartung, S., Buttner, K., and Hopfner, K. P. (2008) Structural biochemistry of a bacterial checkpoint protein reveals diadenylate cyclase activity regulated by DNA recombination intermediates. Mol Cell 30, 167–178

14. Commichau, F. M., Dickmanns, A., Gundlach, J., Ficner, R., and Stülke, J. (2015) A jack of all trades: the multiple roles of the unique essential second messenger cyclic di-AMP. Mol Microbiol 97, 189–204

15. Corrigan, R. M., Abbott, J. C., Burhenne, H., Kaever, V., and Gründling, A. (2011) c-di-AMP is a new second messenger in *Staphylococcus aureus* with a role in controlling cell size and envelope stress. PLoSPathog 7, e1002217

16. Corrigan, R. M., and Gründling, A. (2013) Cyclic di-AMP: another second messenger enters the fray. Nat Rev Microbiol 11, 513–524

17. Mehne, F. M., Gunka, K., Eilers, H., Herzberg, C., Kaever, V., and Stülke, J. (2013) Cyclic di-AMP homeostasis in *Bacillus subtilis*: both lack and high level accumulation of the nucleotide are detrimental for cell growth. J Biol Chem 288, 2004–2017

18. Woodward, J. J., Iavarone, A. T., and Portnoy, D. A. (2010) c-di-AMP secreted by intracellular *Listeria monocytogenes* activates a host type I interferon response. Science 328, 1703–1705

19. Corrigan, R. M., Bellows, L. E., Wood, A., and Gründling, A. (2016) ppGpp negatively impacts ribosome assembly affecting growth and antimicrobial tolerance in Gram-positive bacteria. Proc Natl Acad Sci U S A

20. Bowman, L., Zeden, M. S., Schuster, C. F., Kaever, V., and Gründling, A. (2016) New Insights into the Cyclic Di-adenosine Monophosphate (c-di-AMP) Degradation Pathway and the Requirement of the Cyclic Dinucleotide for Acid Stress Resistance in *Staphylococcus aureus*. J Biol Chem 291, 26970–26986

21. Dengler, V., McCallum, N., Kiefer, P., Christen, P., Patrignani, A., Vorholt, J. A., Berger-Bächi, B., and Senn, M. M. (2013) Mutation in the C-di-AMP cyclase *dacA* affects fitness and resistance of methicillin resistant *Staphylococcus aureus*. PLoS One 8, e73512

22. Gundlach, J., Mehne, F. M., Herzberg, C., Kampf, J., Valerius, O., Kaever, V., and Stülke, J. (2015) An Essential Poison: Synthesis and Degradation of Cyclic Di-AMP in *Bacillus subtilis*. J. Bacteriol 197, 3265–3274

23. Corrigan, R. M., Bowman, L., Willis, A. R., Kaever, V., and Gründling, A. (2015) Cross-talk between two nucleotide-signaling pathways in *Staphylococcus aureus*. J Biol Chem 290, 5826–5839

24. Corrigan, R. M., Campeotto, I., Jeganathan, T., Roelofs, K. G., Lee, V. T., and Gründling, A. (2013) Systematic identification of conserved bacterial c-di-AMP receptor proteins. Proc Natl Acad Sci U S A 110,9084–9089

25. Moscoso, J. A., Schramke, H., Zhang, Y., Tosi, T., Dehbi, A., Jung, K., and Gründling, A. (2015) Binding of Cyclic Di-AMP to the *Staphylococcus aureus* Sensor Kinase KdpD Occurs via the Universal Stress Protein Domain and Downregulates the Expression of the Kdp Potassium Transporter. J Bacteriol 198, 98–110

26. Schuster, C. F., Bellows, L. E., Tosi, T., Campeotto, I., Corrigan, R. M., Freemont, P., and Gründling, A. (2016) The second messenger c-di-AMP inhibits the osmolyte uptake system OpuC in *Staphylococcus aureus*. Sci Signal 9, ra81

27. Campeotto, I., Zhang, Y., Mladenov, M. G., Freemont, P. S., and Gründling, A. (2015) Complex structure and biochemical characterization of the *Staphylococcus aureus* cyclic diadenylate monophosphate (c-di-AMP)-binding protein PstA, the founding member of a new signal transduction protein family. J Biol Chem 290, 2888–2901

28. Chin, K. H., Liang, J. M., Yang, J. G., Shih, M. S., Tu, Z. L., Wang, Y. C., Sun, X. H., Hu, N. J., Liang, Z. X., Dow, J. M., Ryan, R. P., and Chou, S. H. (2015) Structural Insights into the Distinct Binding Mode of Cyclic Di-AMP with SaCpaA_RCK. Biochemistry 54, 4936–4951

29. Kim, H., Youn, S. J., Kim, S. O., Ko, J., Lee, J. O., and Choi, B. S. (2015) Structural Studies of Potassium Transport Protein KtrA Regulator of Conductance of K+ (RCK) C Domain in Complex with Cyclic Diadenosine Monophosphate (c-di-AMP). J Biol Chem 290, 16393–16402

30. Gries, C. M., Bose, J. L., Nuxoll, A. S., Fey, P. D., and Bayles, K. W. (2013) The Ktr potassium transport system in *Staphylococcus aureus* and its role in cell physiology, antimicrobial resistance and pathogenesis. Mol Microbiol 89, 760–773

31. Müller, M., Hopfner, K. P., and Witte, G. (2015) c-di-AMP recognition by *Staphylococcus aureus* PstA. FEBS Lett 589, 45–51

32. Choi, P. H., Sureka, K., Woodward, J. J., and Tong, L. (2015) Molecular basis for the recognition of cyclic-di-AMP by PstA, a PII-like signal transduction protein. Microbiologyopen 4, 361–374

33. Fey, P. D., Endres, J. L., Yajjala, V. K., Widhelm, T. J., Boissy, R. J., Bose, J. L., and Bayles, K. W. (2013) A genetic resource for rapid and comprehensive phenotype screening of nonessential *Staphylococcus aureus* genes. MBio 4, e00537–00512

34. Whiteley, A. T., Pollock, A. J., and Portnoy, D. A. (2015) The PAMP c-di-AMP Is Essential for *Listeria monocytogenes* Growth in Rich but Not Minimal Media due to a Toxic Increase in (p)ppGpp. Cell Host Microbe 17, 788–798

35. Whiteley, A. T., Garelis, N. E., Peterson, B. N., Choi, P. H., Tong, L., Woodward, J. J., and Portnoy, D. A. (2017) c-di-AMP modulates *Listeria monocytogenes* central metabolism to regulate growth, antibiotic resistance and osmoregulation. Mol Microbiol 104, 212–233

36. Halsey, C. R., Lei, S., Wax, J. K., Lehman, M. K., Nuxoll, A. S., Steinke, L., Sadykov, M., Powers, R., and Fey, P. D. (2017) Amino Acid Catabolism in *Staphylococcus aureus* and the Function of Carbon Catabolite Repression. MBio 8

37. Gundlach, J., Herzberg, C., Kaever, V., Gunka, K., Hoffmann, T., Weiss, M., Gibhardt, J., Thurmer, A., Hertel, D., Daniel, R., Bremer, E., Commichau, F. M., and Stülke, J. (2017) Control of potassium homeostasis is an essential function of the second messenger cyclic di-AMP in *Bacillus subtilis*. Sci Signal 10

38. Oppenheimer-Shaanan, Y., Wexselblatt, E., Katzhendler, J., Yavin, E., and Ben-Yehuda, S. (2011) c-di-AMP reports DNA integrity during sporulation in *Bacillus subtilis*. EMBO Rep 12, 594–601

39. Gundlach, J., Herzberg, C., Hertel, D., Thurmer, A., Daniel, R., Link, H., and Stülke, J. (2017) Adaptation of *Bacillus subtilis* to Life at Extreme Potassium Limitation. MBio 8

40. Gundlach, J., Commichau, F. M., and Stülke, J. (2017) Perspective of ions and messengers: an intricate link between potassium, glutamate, and cyclic di-AMP. Curr Genet

41. Bai, Y., Yang, J., Zarrella, T. M., Zhang, Y., Metzger, D. W., and Bai, G. (2014) Cyclic di-AMP impairs potassium uptake mediated by a cyclic di-AMP binding protein in *Streptococcus pneumoniae*. J. Bacteriol 196, 614–623

42. Shapiro, H. M. (2008) Flow cytometry of bacterial membrane potential and permeability. Methods Mol Med 142, 175–186

43. Wetzel, K. J., Bjorge, D., and Schwan, W. R. (2011) Mutational and transcriptional analyses of the *Staphylococcus aureus* low-affinity proline transporter OpuD during in vitro growth and infection of murine tissues. FEMS Immunol Med Microbiol 61, 346–355

44. Price-Whelan, A., Poon, C. K., Benson, M. A., Eidem, T. T., Roux, C. M., Boyd, J. M., Dunman, P. M., Torres, V. J., and Krulwich, T. A. (2013) Transcriptional profiling of *Staphylococcus aureus* during growth in 2 M NaCl leads to clarification of physiological roles for Kdp and Ktr K+ uptake systems. MBio 4

45. Choi, S., Jung, J., Jeon, C. O., and Park, W. (2014) Comparative genomic and transcriptomic analyses of NaCl-tolerant *Staphylococcus* sp. OJ82 isolated from fermented seafood. Appl Microbiol Biotechnol 98, 807–822

46. Kafala, B., and Sasarman, A. (1994) Cloning and sequence analysis of the *hemB* gene of *Staphylococcus aureus*. Can J Microbiol 40, 651–657

47. von Eiff, C., Heilmann, C., Proctor, R. A., Woltz, C., Peters, G., and Götz, F. (1997) A site-directed *Staphylococcus aureus hemB* mutant is a small-colony variant which persists intracellularly. J Bacteriol 179, 4706–4712

48. Imlay, J. A. (2003) Pathways of oxidative damage. Annu Rev Microbiol 57, 395–418

49. Li, G. Q., Quan, F., Qu, T., Lu, J., Chen, S. L., Cui, L. Y., Guo, D. W., and Wang, Y. C. (2015) Sublethal vancomycin-induced ROS mediating antibiotic resistance in *Staphylococcus aureus*. Biosci Rep 35

50. Tavares, A. F., Teixeira, M., Romao, C. C., Seixas, J. D., Nobre, L. S., and Saraiva, L. M. (2011) Reactive oxygen species mediate bactericidal killing elicited by carbon monoxide-releasing molecules. J Biol Chem 286, 26708–26717

51. Griffiths, J. M., and O’Neill, A. J. (2012) Loss of function of the *gdpP* protein leads to joint beta-lactam/glycopeptide tolerance in *Staphylococcus aureus*. Antimicrob Agents Chemother 56, 579–581

52. Pozzi, C., Waters, E. M., Rudkin, J. K., Schaeffer, C. R., Lohan, A. J., Tong, P., Loftus, B. J., Pier, G. B., Fey, P. D., Massey, R. C., and O’Gara, J. P. (2012) Methicillin resistance alters the biofilm phenotype and attenuates virulence in *Staphylococcus aureus* device-associated infections. PLoS Pathog 8, e1002626

53. Angelidis, A. S., and Smith, G. M. (2003) Three transporters mediate uptake of glycine betaine and carnitine by *Listeria monocytogenes* in response to hyperosmotic stress. Appl Environ Microbiol 69, 1013–1022

54. Ko, R., and Smith, L. T. (1999) Identification of an ATP-driven, osmoregulated glycine betaine transport system in *Listeria monocytogenes*. Appl Environ Microbiol 65, 4040–4048

55. Huynh, T. N., Choi, P. H., Sureka, K., Ledvina, H. E., Campillo, J., Tong, L., and Woodward, J. J. (2016) Cyclic di-AMP targets the cystathionine beta-synthase domain of the osmolyte transporter OpuC. Mol Microbiol

56. Sleator, R. D., Gahan, C. G., Abee, T., and Hill, C. (1999) Identification and disruption of BetL, a secondary glycine betaine transport system linked to the salt tolerance of *Listeria monocytogenes* LO28. Appl Environ Microbiol 65, 2078–2083

57. Hiron, A., Borezee-Durant, E., Piard, J. C., and Juillard, V. (2007) Only one of four oligopeptide transport systems mediates nitrogen nutrition in *Staphylococcus aureus*. J Bacteriol 189, 5119–5129

58. Yu, D., Pi, B., Yu, M., Wang, Y., Ruan, Z., Feng, Y., and Yu, Y. (2014) Diversity and evolution of oligopeptide permease systems in *Staphylococcal* species. Genomics 104, 8–13

59. Seggewiss, J., Becker, K., Kotte, O., Eisenacher, M., Yazdi, M. R., Fischer, A., McNamara, P., Al Laham, N., Proctor, R., Peters, G., Heinemann, M., and von Eiff, C. (2006) Reporter metabolite analysis of transcriptional profiles of a *Staphylococcus aureus* strain with normal phenotype and its isogenic *hemB* mutant displaying the small-colony-variant phenotype. J Bacteriol 188, 7765–7777

60. Kohler, C., von Eiff, C., Peters, G., Proctor, R. A., Hecker, M., and Engelmann, S. (2003) Physiological characterization of a heme-deficient mutant of *Staphylococcus aureus* by a proteomic approach. J Bacteriol 185, 6928–6937

61. Kohler, C., von Eiff, C., Liebeke, M., McNamara, P. J., Lalk, M., Proctor, R. A., Hecker, M., and Engelmann, S. (2008) A defect in menadione biosynthesis induces global changes in gene expression in *Staphylococcus aureus*. J Bacteriol 190, 6351–6364

62. Zamboni, N., and Sauer, U. (2003) Knockout of the high-coupling cytochrome aa3 oxidase reduces TCA cycle fluxes in *Bacillus subtilis*. FEMS Microbiol Lett 226, 121–126

63. Commichau, F. M., Gibhardt, J., Halbedel, S., Gundlach, J., and Stülke, J. (2017) A Delicate Connection: c-di-AMP Affects Cell Integrity by Controlling Osmolyte Transport. Trends Microbiol

64. Rudin, L., Sjostrom, J. E., Lindberg, M., and Philipson, L. (1974) Factors affecting competence for transformation in *Staphylococcus aureus*. J Bacteriol 118, 155–164

65. Townsend, D. E., and Wilkinson, B. J. (1992) Proline transport in *Staphylococcus aureus:* a high-affinity system and a low-affinity system involved in osmoregulation. J Bacteriol 174, 2702–2710

66. Monk, I. R., Shah, I. M., Xu, M., Tan, M. W., and Foster, T. J. (2012) Transforming the untransformable: application of direct transformation to manipulate genetically *Staphylococcus aureus* and *Staphylococcus epidermidis*. MBio 3

67. Behrends, V., Bell, T. J., Liebeke, M., Cordes-Blauert, A., Ashraf, S. N., Nair, C., Zlosnik, J. E., Williams, H. D., and Bundy, J. G. (2013) Metabolite profiling to characterize disease-related bacteria: gluconate excretion by *Pseudomonas aeruginosa* mutants and clinical isolates from cystic fibrosis patients. J Biol Chem 288, 15098–15109

68. Mayer, S., Steffen, W., Steuber, J., and Gotz, F. (2015) The *Staphylococcus aureus* NuoL-like protein MpsA contributes to the generation of membrane potential. J Bacteriol 197, 794–806

69. Charpentier, E., Anton, A. I., Barry, P., Alfonso, B., Fang, Y., and Novick, R. P. (2004) Novel cassette-based shuttle vector system for gram-positive bacteria. Appl Environ Microbiol 70, 6076–6085

70. Gründling, A., and Schneewind, O. (2007) Genes required for glycolipid synthesis and lipoteichoic acid anchoring in *Staphylococcus aureus*. J Bacteriol 189, 2521–2530

71. Monk, I. R., Tree, J. J., Howden, B. P., Stinear, T. P., and Foster, T. J. (2015) Complete Bypass of Restriction Systems for Major *Staphylococcus aureus* Lineages. MBio 6, e00308–00315

72. Horsburgh, M. J., Aish, J. L., White, I. J., Shaw, L., Lithgow, J. K., and Foster, S. J. (2002) sigmaB modulates virulence determinant expression and stress resistance: characterization of a functional *rsbU* strain derived from *Staphylococcus aureus* 8325-4. J Bacteriol 184, 5457–5467

73. Cosgrove, K., Coutts, G., Jonsson, I. M., Tarkowski, A., Kokai-Kun, J. F., Mond, J. J., and Foster, S. J. (2007) Catalase (KatA) and alkyl hydroperoxide reductase (AhpC) have compensatory roles in peroxide stress resistance and are required for survival, persistence, and nasal colonization in *Staphylococcus aureus*. J Bacteriol 189, 1025–1035

74. Pader, V., James, E. H., Painter, K. L., Wigneshweraraj, S., and Edwards, A. M. (2014) The Agr quorum-sensing system regulates fibronectin binding but not hemolysis in the absence of a functional electron transport chain. Infect Immun 82, 4337–4347

75. Boles, B. R., Thoendel, M., Roth, A. J., and Horswill, A. R. (2010) Identification of genes involved in polysaccharide-independent *Staphylococcus aureus* biofilm formation. PLoS One 5, e10146

